# The Neural Architecture of Theory-based Reinforcement Learning

**DOI:** 10.1101/2022.06.14.496001

**Authors:** Momchil S. Tomov, Pedro A. Tsividis, Thomas Pouncy, Joshua B. Tenenbaum, Samuel J. Gershman

## Abstract

Humans learn internal models of the environment that support efficient planning and flexible generalization in complex, real-world domains. Yet it remains unclear how such internal models are represented and learned in the brain. We approach this question within the framework of theory-based reinforcement learning, a strong form of model-based reinforcement learning in which the model is an intuitive theory – a rich, abstract, causal model of the environment built on a natural ontology of physical objects, intentional agents, relations, and goals. We used a theory-based reinforcement learning model to analyze brain data from human participants learning to play different Atari-style video games while undergoing functional MRI. Theories inferred by the theory-based model explained the signal in inferior frontal gyrus and other prefrontal areas better than several alternative models. Brain activity increased in response to theory update events in inferior frontal gyrus, occipital cortex, and fusiform gyrus, with separate learning signals for different theory components. This corresponded with a transient strengthening of theory representations in those regions. Finally, the effective connectivity pattern during theory updating suggests that information flows top-down from theory-coding regions in the prefrontal cortex to theory updating regions in occipital and temporal cortex. These results are consistent with a neural architecture in which top-down theory representations originating in prefrontal regions shape sensory predictions in visual areas, where factorized theory prediction errors are computed and in turn trigger bottom-up updates of the theory. This initial sketch provides a foundation for understanding of the neural representations and computations that support efficient theory-based reinforcement learning in complex, naturalistic environments.

## 1. Introduction

Reinforcement learning (RL) is a normative framework prescribing how agents ought to act in order to maximize rewards in the environment (Sutton and Barto, 2018). In the field of artificial intelligence, RL has allowed artificial agents to reach and surpass human-level performance across a variety of domains previously beyond the capabilities of computers (Mnih et al., 2015; Silver et al., 2017, 2018; Schrittwieser et al., 2020). In the fields of psychology and neuroscience, RL has offered a compelling account of behavioral and brain data across a number of species and experimental paradigms (Schultz et al., 1997; Daw et al., 2006; Niv, 2009; Cross et al., 2021). Most of this work has focused on model-free RL, a kind of RL in which the agent directly learns a mapping from different states in the environment to actions and/or values. Model-based RL, on the other hand, posits that the agent learns an internal model of the environment which is used to simulate the outcomes of different actions. Behavioral and neural studies have found evidence for both kinds of RL (Gläscher et al., 2010; Daw et al., 2011; Lee et al., 2014; Kool et al., 2018), yet model-based RL has received relatively less attention and is often studied using simple toy environments with small state spaces. This is largely owing to the relative scarcity of powerful model-based RL algorithms capable of matching human learning in complex domains (Tsividis et al., 2017), leaving open the question of what the “model” in model-based RL is and how it is learned and represented by the brain.

One possible answer from cognitive science is theory-based RL (Pouncy et al., 2021; Tsividis et al., 2021; Pouncy and Gershman, 2022), a strong form of model-based RL in which the model is an intuitive theory—an abstract causal model of world dynamics rooted in core cognitive concepts such as physical objects, intentional agents, relations, and goals (Figure 1). Building on findings in developmental psychology, theory-based RL posits that the agent learns the theory from experience using probabilistic inference and uses it together with an internal simulator to predict and evaluate the outcomes of different action sequences generated by an internal planner. Theory-based RL has captured patterns of human learning (Pouncy and Gershman, 2022; Tsividis et al., 2021), exploration (Tsividis et al., 2021), and generalization (Pouncy et al., 2021) in complex domains where model-free and simpler model-based RL approaches fail or learn rather differently. This has provided strong support for theory-based RL as a concrete realization of human model-based RL. Building on this work, our study aims to identify brain regions involved in theory-based RL and how they map to its constituent processes. To achieve this, we used a particular formalization of theory-based RL (Tsividis et al., 2021) to analyze functional magnetic resonance imaging (fMRI) data collected from human participants while they learned to play Atari-style games designed to mirror some of the richness and complexity of real-world tasks. Our analyses revealed evidence that theory representations in inferior frontal gyrus and other prefrontal regions are activated and updated in response to theory prediction errors—discrepancies between theoretical predictions and actual observations—which are in turn computed in occipital and ventral stream regions such as the fusiform gyrus. We also found evidence that, much like in our theory-based RL model, theory updating in the brain is factorized into updating of objects, relations, and goals, suggesting key differences between these cognitive components. Finally, analyses of effective connectivity suggest that theory inference involves both feedforward and feedback processing reminiscent of hierarchical predictive coding (Rao and Ballard, 1999; Friston, 2005). Together these results present the first direct evidence for theory-based RL in the brain and establish a foundation for understanding its underlying neural processes.

**Figure 1:**
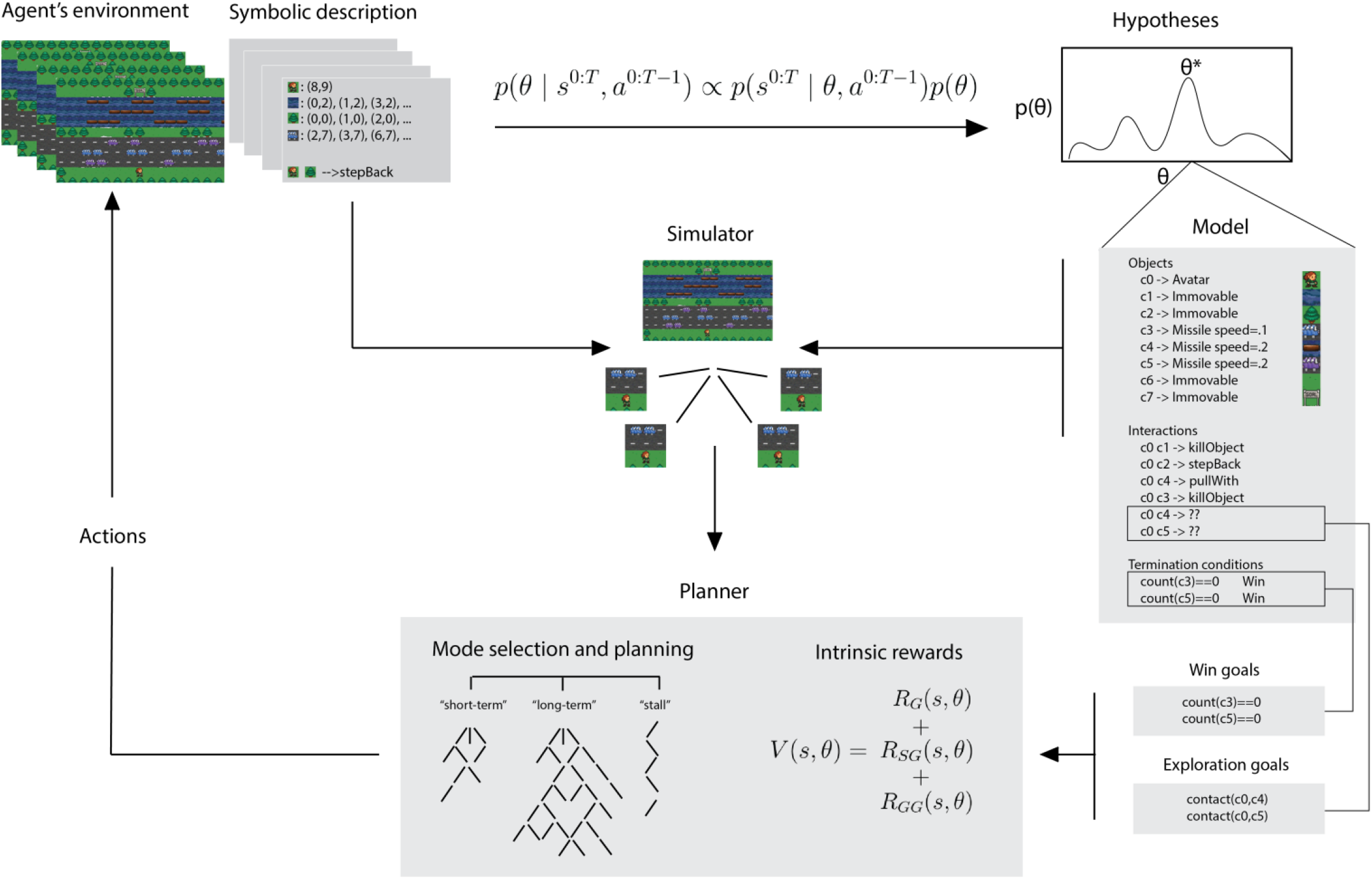
EMPA architecture. Symbolic descriptions of game frames are fed to an inference engine which updates the most likely theory, *θ*^*^, using an approximation of Bayesian inference. The theory consists of objects (sprites), relations (interactions), and goals (termination conditions). Exploitative (win) and exploratory goals based on the theory are fed to a planner which uses a theory-based internal simulator and an intrinsic reward function to search for rewarding action sequences. The agent then takes actions in the environment according to the best plan. Reused with permission from Tsividis et al. (2021).

## 2. Results

We scanned 32 human participants using fMRI while they played six Atari-style games (Figure 2A). Each game had nine levels of increasing complexity and had to be learned from experience without any visual hints or prior information about the rules. For data analysis purposes, games were interleaved and balanced across pairs of runs (Figure 2B).

**Figure 2:**
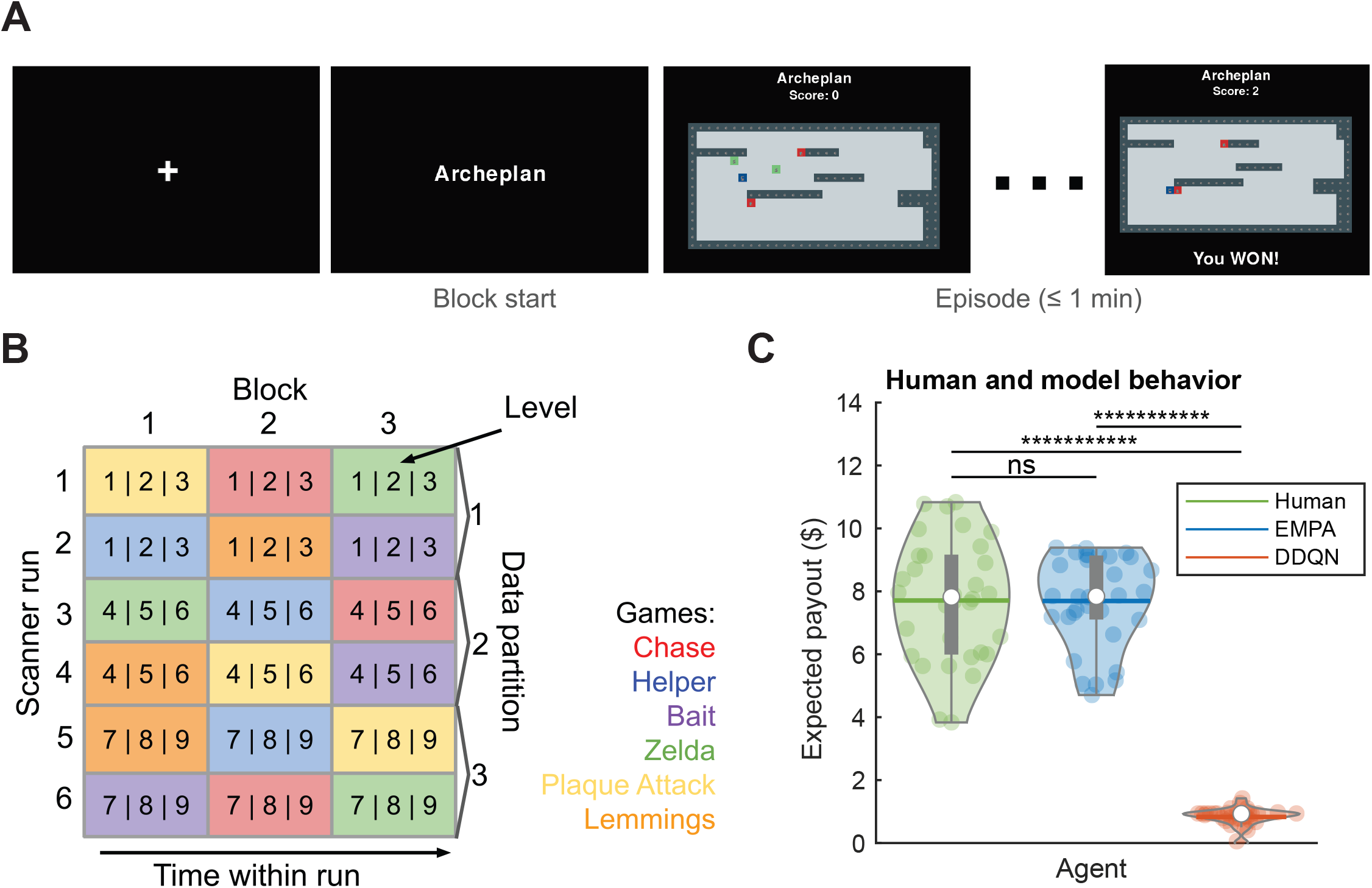
Study design and behavioral results. A. Participants played different Atari-style games. The game names, colors, and textures shown were randomized for each participant and were unrelated to the game rules (but were consistent across different levels of the same game for each participant). B. Example scan session for single participant. Runs were paired into balanced data partitions, with blocks shuffled within each partition. C. Behavioral results from participants (green) and generative play by EMPA (blue) and pretrained DDQN (red). Each colored dot represents a single real or simulated participant, respectively. White dots depict medians, box plots depict upper and lower quartiles, horizontal lines across kernel density estimates depict means. ns - not significant, ********** - *p <* 10^*−*10^ (two-sided Wilcoxon rank sum test).

As a particular instantiation of theory-based RL, we used the Explore, Model, Plan Agent (EMPA; Figure 1) proposed by Tsividis et al. (2021). Theories are formalized as symbolic, probabilistic program-like descriptions of game dynamics that specify the different object kinds, the outcomes of interactions between them, and the win/loss conditions. EMPA performs Bayesian inference over the space of theories and uses the most likely theory to run internal simulations and search for rewarding action sequences. Tsividis et al. (2021) showed that EMPA exhibits human-level learning efficiency in a large suite of Atari-style games, including those used in our study. They also showed that EMPA exhibits human-like object-oriented exploratory behaviors. In contrast, model-free RL agents failed on both counts, learning orders of magnitude more slowly and exploring much more randomly than humans. Consistent with these results, we found that EMPA performed similarly to our participants (Figure 2C; no significant difference, two-sided Wilcoxon rank sum test based on simulated and actual expected bonus payouts), while both humans and EMPA performed significantly better than a pretrained deep RL network, the double DQN (DDQN; p < 10^−10^), a powerful model-free RL algorithm (van Hasselt et al., 2016), variants of which have been put forward as accounts of human model-free RL in complex domains (Cross et al., 2021).

## 3. Theory representations in prefrontal cortex

The central component of theory-based RL is the theory that the agent continuously infers from experience. To identify brain regions representing the inferred theory, we replayed each participant’s gameplay through EMPA and used the inferred theory sequences to predict the blood-oxygen-level-dependent (BOLD) signal in each voxel using a linear encoding model fit with Gaussian process (GP) regression (Figure 3A, Figure S3A,C), a generalization of the more commonly used ridge regression (Williams and Rasmussen, 2006). In order to embed the symbolic theories in a vector space for the encoding model, we used holographic reduced representations (HRRs; Plate, 1995), a method for encoding complex compositional structure in distributed form. Similarly to previous work (Schrimpf et al., 2021; Cross et al., 2021), we correlated the cross-validated predicted and actual BOLD timecourses; we then Fisher z-transformed the resulting Pearson correlation coefficients to compute a predictivity score for each voxel. Predictivity scores were aggregated across voxels using t-tests. The resulting group level t-maps were thresholded at *p* < 0.001 and whole-brain cluster family-wise error (FWE) corrected at *α* = 0.05. This revealed significant predictivity scores across a distributed bilateral network of regions (Figure 3B, Table S1). In prefrontal cortex, we found bilateral clusters in inferior frontal gyrus, as well as unilateral clusters in middle and superior frontal gyrus and the supplementary motor area. In posterior areas, we found a large bilateral cluster starting from early visual regions in occipital cortex, extending into higher visual regions and then further into the ventral and dorsal streams, including fusiform gyrus and middle temporal gyrus in temporal cortex and inferior parietal gyrus and angular gyrus in parietal cortex.

**Figure 3:**
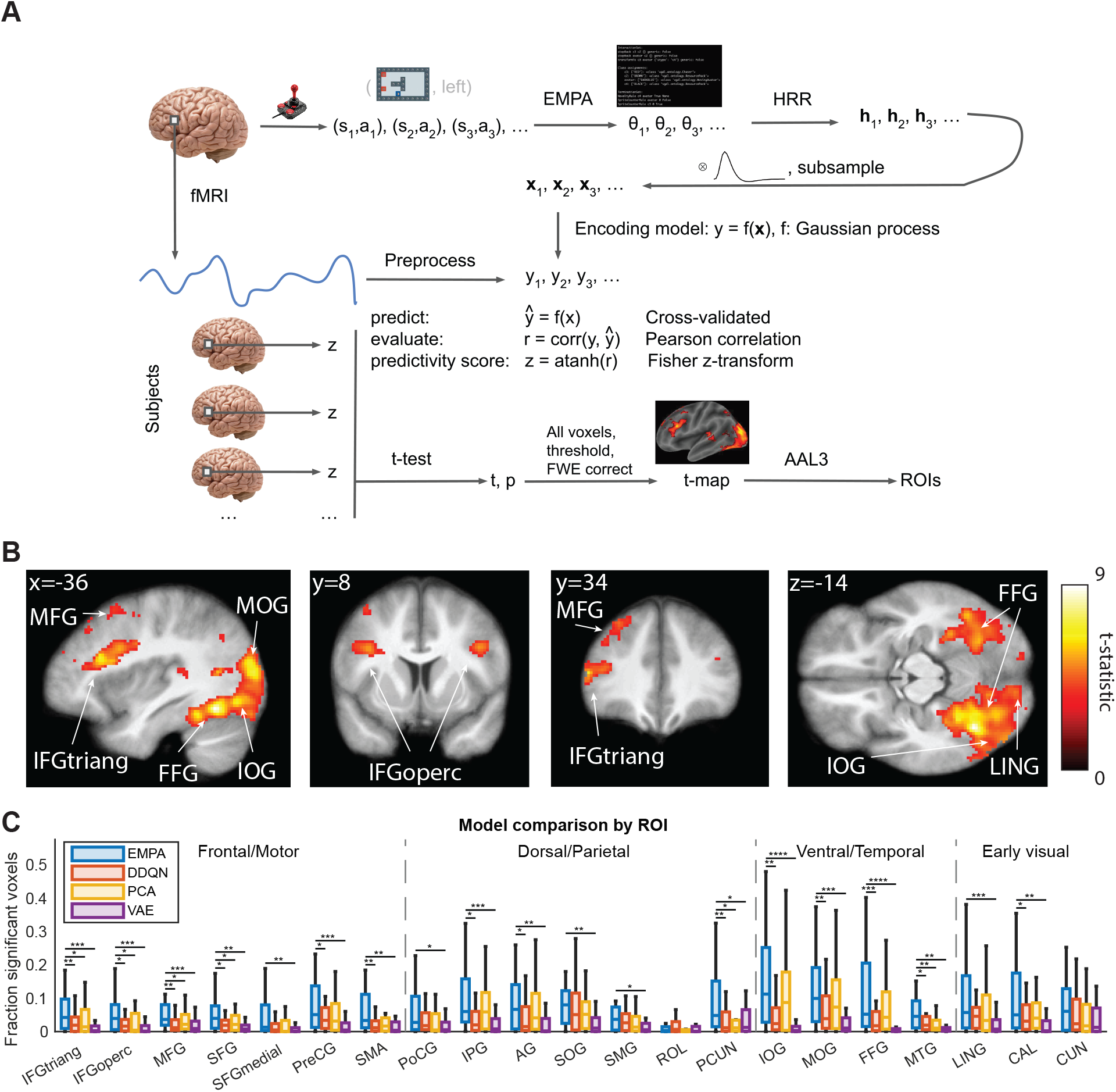
Theory representations map to regions in prefrontal cortex and ventral/dorsal streams. A. Encoding model analysis pipeline. State-action sequences ((*a*_1_, *s*_1_), (*s*_2_, *a*_2_), (*s*_3_, *a*_3_), …) from human gameplay were replayed through EMPA. Inferred theory sequences (*θ*_1_, *θ*_2_, *θ*_3_, …) were embedded in a vector space, convolved with the hemodynamic response function, and subsampled to get feature vectors (**x**_1_, **x**_2_, **x**_3_, …). Preprocessed BOLD signal from each voxel (*y*_1_, *y*_2_, *y*_3_, …) was regressed onto feature vectors using GP regression. Resulting predictivity scores *z* were aggregated across participants using two-sided t-tests. Resulting t-maps were thresholded at *p <* 0.001 and whole-brain cluster FWE corrected at *α* = 0.05. Analogous analyses were performed with control models (DDQN, PCA, VAE). B. Group-level t-maps from A. ROIs are noted as IFGtriang, IFGoperc (inferior frontal gyrus, triangular and opercular parts), MFG, SFG (middle and superior frontal gyri), PreCG (precentral gyrus), SMA (supplementary motor area), PoCG (postcentral gyrus), IPG (inferior parietal gyrus), AG (angular gyrus), SMG (supramarginal gyrus), ROL (rolandic operculum), PCUN (precuneus), IOG, MOG, SOG (inferior, middle, superior orbital gyri), FFG (fusiform gyrus), MTG (middle temporal gyrus), LING (lingual gyrus), CAL (calcarine fissure), CUN (cuneus). C. Fraction of voxels with significant correlation (*α* = 0.05) between predicted and actual BOLD in anatomical ROIs. Medians with boxes representing top and bottom quartiles and whiskers representing data range, excluding outliers (outliers plotted in Figure S3A and included in all statistical tests). * - *p <* 0.05, ** - *p <* 0.01, *** - *p <* 0.001, **** - *p <* 0.0001 (two-sided Wilcoxon signed rank tests).

We performed the same analysis using three control models (Cross et al., 2021): DDQN agents pretrained on corresponding games to control for model-free RL representations (also used as a behavioral control in Tsividis et al., 2021), principal component analysis (PCA) to control for low-level visual features (Olshausen and Field, 1996; Chang and Tsao, 2017), and a variational autoencoder (VAE) to account for high-level visual and state features (Mohamed and Jimenez Rezende, 2015; Watter et al., 2015; Higgins et al., 2017). Similarly to Cross et al. (2021), we compared models using *a posteriori* bilateral anatomical regions of interest (ROIs) based on cross-referencing the t-maps from all models with the automated anatomical labelling atlas (AAL3 atlas; Rolls et al., 2020). We compared models separately in each ROI based on the fraction of voxels with a significant correlation (*α* = 0.05) between predicted and actual BOLD signal. In prefrontal regions, EMPA largely outperformed all three control models, specifically in the triangular and opercular parts of inferior frontal gyrus, as well as in the middle and superior frontal gyri. Additionally, EMPA outperformed all three control models in precuneus and middle temporal gyrus. This suggests that the effects of theory representation in those regions are not simply due to visual or model-free RL confounds. We also repeated the analysis using different components of the EMPA theory – objects, relations, and goals – but did not find any systematic differences (Figure S3B,D).

## 4. Theory update signals in inferior frontal gyrus, occipital gyri, and fusiform gyrus

After identifying regions representing the inferred theory, we next sought to identify brain regions involved in theory inference. Based on our previous work (Tomov et al., 2018), we reasoned that such regions might show greater activity during theory updating, reflecting the temporary increase in computational demands. Since theory updates are triggered by surprising events which violate theoretical predictions, such an increase in neural activity could also be interpreted as a kind of theory prediction error. We used a general linear model (GLM) with impulse regressors at theory update events – frames at which EMPA switched from one most likely theory to another based on the participant’s gameplay (Figure 4A, Table S2). The group-level contrast for theory updating (Figure 4B, Table S3; thresholded at p < 0.001 and whole-brain cluster FWE corrected at *α* = 0.05) revealed a distributed bilateral network of regions that largely overlapped with the regions from our theory representation analysis. Most notably, in prefrontal cortex, we found bilateral clusters in inferior frontal gyrus, in addition to unilateral clusters in superior frontal gyrus, orbital frontal cortex, and the supplementary motor area. We also found a large bilateral posterior cluster covering early and late visual regions in occipital cortex, extending into angular gyrus and precuneus in the dorsal stream, and extending into fusiform gyrus in the ventral stream.

**Figure 4:**
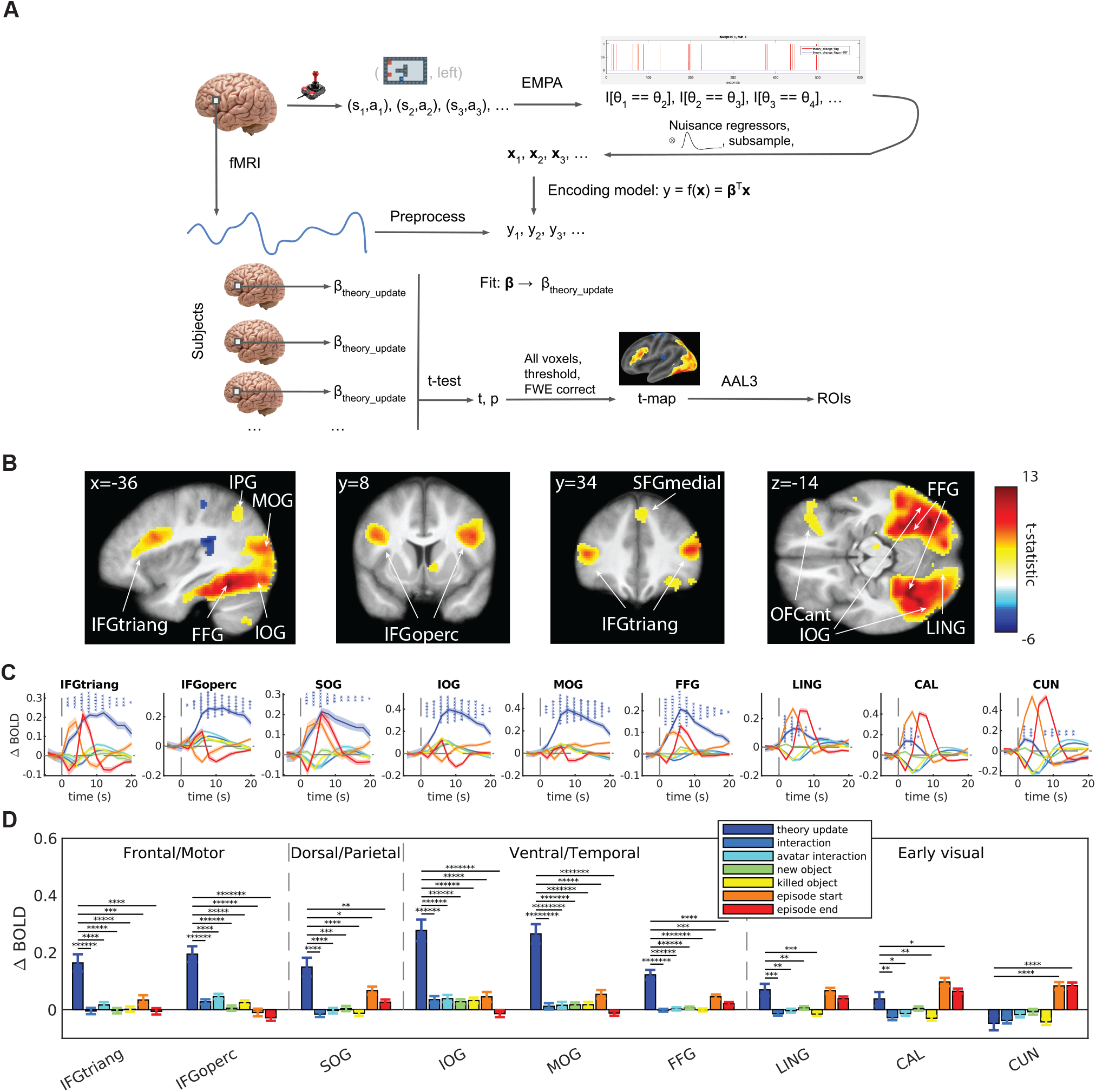
Theory learning signals in prefrontal cortex and ventral/dorsal streams. A. GLM analysis pipeline. Similarly to Figure 3A, frame-by-frame state-action sequences ((*a*_1_, *s*_1_), (*s*_2_, *a*_2_), (*s*_3_, *a*_3_), …) from human gameplay were replayed through EMPA. Corresponding theory update sequences (*I*[*θ*_1_ ≡ *θ*_2_], *I*[*θ*_2_ ≡ *θ*_3_], *I*[*θ*_3_ ≡ *θ*_4_], …) from EMPA were entered as regressors in a GLM. Resulting theory update beta estimates (*β*_theory update_) for individual voxels aggregated across participants using two-sided t-tests. Resulting t-maps thresholded at *p <* 0.001 and whole-brain cluster FWE corrected at *α* = 0.05. B. Group-level t-maps from GLM analysis in A. ROIs noted as OFCant (anterior orbital gyrus), SFGmedial (superior frontal gyrus, medial), and the rest as in Figure 3B. C. Peri-event time histograms showing the average change in BOLD signal following theory updates and different control events in ROIs with significant *β*_theory update_. Colored fringes depict error bars (s.e.m.) across participants. Stars indicate significance for theory updates for each time point. * - *p <* 0.05, ** - *p <* 0.01, *** - *p <* 0.001, **** - *p <* 0.0001, ***** - *p <* 0.00001, ****** - *p <* 10^*−*6^ (two-sided t-tests). D. Change in BOLD signal from C averaged over 20 s following corresponding event. Error bars depict s.e.m. across participants. Significance notation as in C (paired t-tests).

To control for potential confounds, we included a number of nuisance regressors in the GLM for events of non-interest, including visual changes, key presses, and game events relevant for theory updating (Table S2). A confirmatory analysis using anatomical ROIs from the theory updating contrast revealed that some nuisance regressors also show a significant effect (Figure S4). To directly compare the neural responses to different event types, we generated peri-event time histograms (PETHs) from the baseline-adjusted BOLD signal following theory updates and other control events (Figure 4C, D) in bilateral anatomical ROIs with a significant theory update effect (Figure S4). We found that, in contrast to other control events, the increase in BOLD signal was larger and more sustained after theory updates in inferior frontal gyrus, all three occipital gyri, and fusiform gyrus (two-sided t-tests in Figure 4C, paired t-tests in Figure 4D). These results suggest that these regions respond specifically to theory updating, pointing to their potential involvement in computing theory prediction errors – discrepancies between the perceived world state and the predicted world state based on the theory – or in performing the theory update computation in response to such errors. It is also noteworthy that these regions also appear in the theory representation brain maps (Figure 3B), with inferior frontal gyrus specifically representing the learned theory (Figure 3C).

## 5. Separate update signals for different theory components

The EMPA theory consists of three components: a set of object types and their physical and/or intentional properties (since they could be other agents), a set of relations between objects describing the outcomes of object-object interactions, and a set of goals that the agent pursues. For tractability, EMPA factorizes theory inference into separate inference processes for objects, relations, and goals (Tsividis et al., 2021). However, the theory update GLM described above does not distinguish between updates for separate theory components. Rather, theory update events occur when either objects, relations, or goals are updated (Figure 5A, top). When we repeated the PETH analysis described above for individual theory component updates, we found that some regions respond differentially to different component updates (Figure S5). This led us to hypothesize that the brain might factorize theory learning similarly to EMPA.

**Figure 5:**
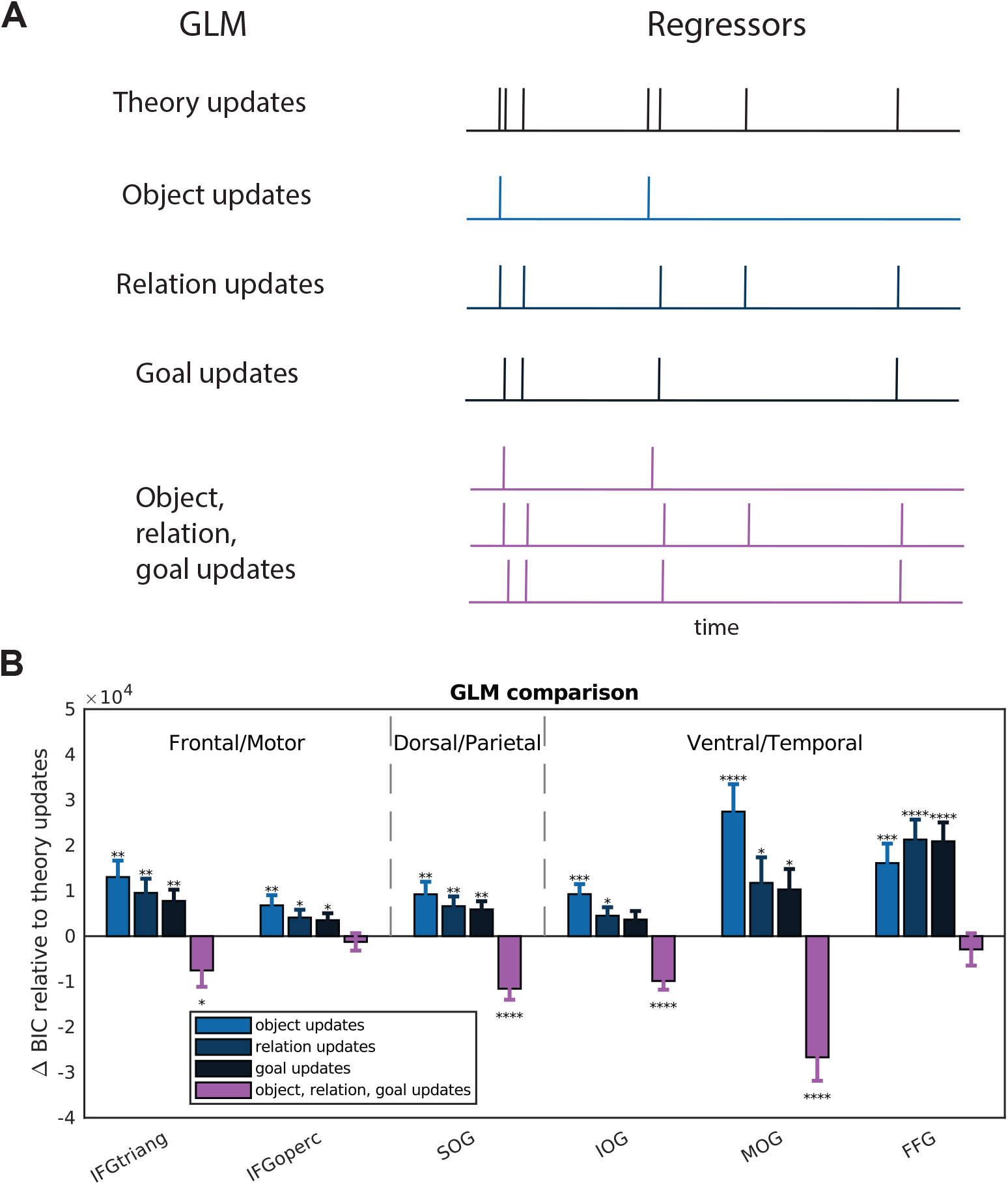
Separate update signals for different theory components. A. Illustration of GLMs with impulse regressors for unified theory updates (top GLM; same as in Figure 4), single component updates (middle three GLMs), and separate updates for all three components (bottom GLM). B. GLM comparison in ROIs showing a significant increase in BOLD signal for all three theory components (Figure S5B). ROIs noted as in Figure 3B. Bars denote GLM BICs relative to theory update GLM BIC. Error bars denote s.e.m. across participants. * - *p <* 0.05, ** - *p <* 0.01, *** - *p <* 0.001, **** - *p <* 0.0001 (two-sided t-tests). BIC, Bayesian information criterion.

To investigate this hypothesis, we fit a GLM in which theory updating was split into three separate regressors for object, relation, and goal updates (Figure 5A, bottom). We additionally fit three control GLMs, each with a single component update (Figure 5A, middle). We compared GLMs using random effects Bayesian model selection (Rigoux et al., 2014) in the ROIs showing a significant BOLD increase in response to all three individual component updates (Figure S5B). We found that the GLM with separate component updates best explains the BOLD signal in inferior frontal gyrus and all three occipital gyri (Figure 5B, Table 1). This suggests that, similarly to EMPA, the brain also performs a factorized theory update.

**Table 1:** GLM comparison results. PXP, protected exceedance probability. IFG, inferior frontal gyrus.

|  | GLM PXP |  |  |  |  |
| --- | --- | --- | --- | --- | --- |
| AAL3 Region | Theory updates | Object updates | Relation updates | Goal updates | Object, relation, goal updates |
| IFG pars triangularis | < 0.0001 | < 0.0001 | < 0.0001 | < 0.0001 | 0.9998 |
| IFG pars opercularis | 0.1717 | 0.1717 | 0.1586 | 0.1649 | 0.3328 |
| Superior occipital gyrus | 0.0004 | < 0.0001 | < 0.0001 | < 0.0001 | 0.99953 |
| Inferior occipital gyrus | < 0.0001 | < 0.0001 | < 0.0001 | < 0.0001 | 0.99998 |
| Middle occipital gyrus | < 0.0001 | < 0.0001 | < 0.0001 | < 0.0001 | 0.99996 |
| Fusiform gyrus | 0.7140 | 0.0097 | 0.0004 | 0.0004 | 0.2756 |

## 6. Theory representations activated during updating

The overlap (Figure 6A) between the brain regions representing the theory (Figure 3) and the brain regions responding to theory updating (Figure 4) was somewhat surprising. *A priori*, these regions do not necessarily have to be the same: one analysis looks for regions consistently representing the theory, without any increase in activity around change points, while the other analysis looks for regions with increased activity at theory change points, without regard for the content of the theory itself. This led us to hypothesize that the two are related. Specifically, we conjectured that theory representations are preferentially activated during theory updating, akin to being “loaded” into working memory for the necessary computation.

**Figure 6:**
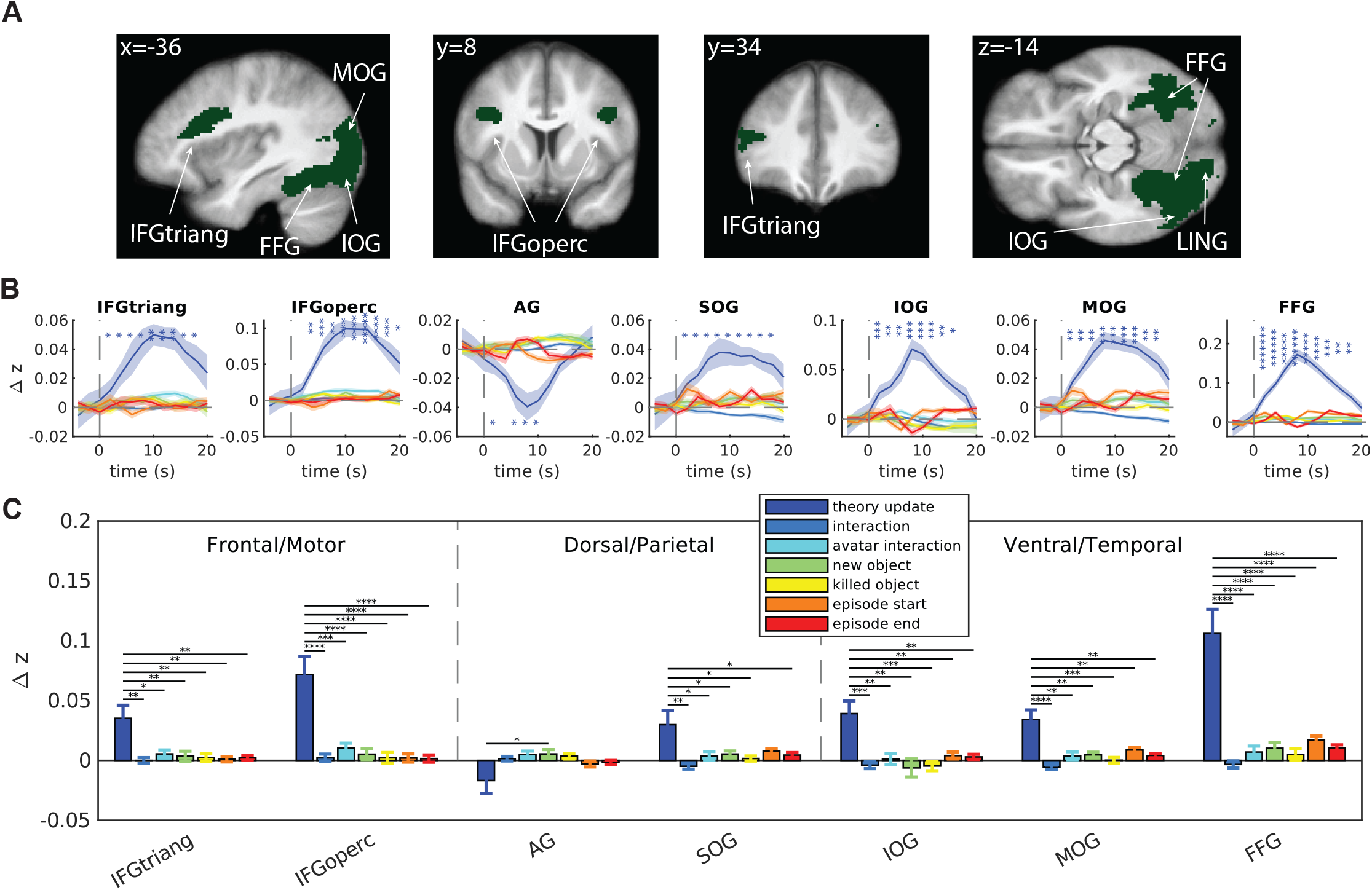
Theory representations activated during updating. A. Overlap between significant clusters for theory representations (Figure 3B) and theory updating (Figure 4B). ROIs noted as in Figure 3B. B. Peri-event time histograms showing the average change in predictivity score following theory updates and different control events in the overlapping ROIs. Notation as in Figure 4C. Note that, in contrast to Figure 4C, the y-axis is Δ*z*, which quantifies how well an encoding model based on theory representations can predict instantaneous patterns of brain activity after a theory update, compared to before the update. C. Change in predictivity score from B averaged over 20 s following corresponding event. Notation as in Figure 4D.

To investigate this hypothesis, we plotted PETHs of the baseline-adjusted predictivity time-course from the encoding model (Figure 3A) following theory updates and other control events in the ROIs from the overlap. This shows, at each time point after the event, how well the pattern of BOLD activity can be predicted based on the inferred theory, compared to immediately before the event. We found a significant sustained increase in predictivity after theory updates in inferior frontal gyrus (triangular and opercular parts), all three occipital gyri (inferior, middle, superior), and fusiform gyrus (Figure 6B; two-sided t-tests). Furthermore, the magnitude of this increase was significantly greater for theory updates compared to other events (Figure 6C; paired t-tests), suggesting that theory representations are activated in these regions specifically during theory updating.

To investigate whether this effect varies between individual theory components, we repeated this analysis for separate component updates using the corresponding encoding models fit for objects, relations, or goals only. We found that most regions did not show a significant difference (Figure S6), with the exception of fusiform gyrus in which object representations were activated after object updates more strongly compared to relation and goal representations during their respective updates (p < 0.001, Bonferroni corrected), suggesting a specific role for fusiform gyrus in object updating.

## 7. Effective connectivity during theory updating is consistent with predictive coding

Having identified brain regions involved in theory representation (Figure 3), theory updating (Figure 4, Figure 5), and the dynamic interplay between these processes (Figure 6), we finally sought to characterize the pattern of information flow between these regions. Using a beta series GLM (Poldrack et al., 2011), we extracted estimates of instantaneous neural activity during theory update events from ROIs that showed a significant effect in the previous analyses. We additionally extracted estimates from visual and motor ROIs in order to include potential inputs and outputs to and from the theory coding and updating regions. We entered the resulting estimates into the IMaGES algorithm (Ramsey et al., 2010; Poldrack et al., 2011) from the `TETRAD` software package for causal modeling (Scheines et al., 1998), which greedily searches the space of effective connectivity patterns for the one that best fits the data. Our hypothesis was that, during theory updating, information would flow in a bottom-up fashion, from early visual regions through theory updating regions in occipital and temporal cortex to theory coding regions in prefrontal cortex, where the updated theory is putatively stored.

To the contrary, we found the opposite pattern, with information flowing in a top-down fashion, from prefrontal theory coding regions to theory updating regions in occipital and temporal cortex to early visual regions (Figure 7A). When we repeated the same analysis, except using neural activity 2 s after theory updates, we found a bottom-up pattern consistent with our prior expectations (Figure 7B). These findings are consistent with a predictive coding interpretation: information about the brain’s internal model of the world (in our case, the theory) is flowing top-down from higher areas in prefrontal cortex, shaping sensory predictions in lower visual areas; when an inconsistency between predictions and observations is detected, this results in a theory prediction error which triggers a theory update, reversing the flow of information so that the new sensory data can be used to update the theory in the higher regions.

**Figure 7:**
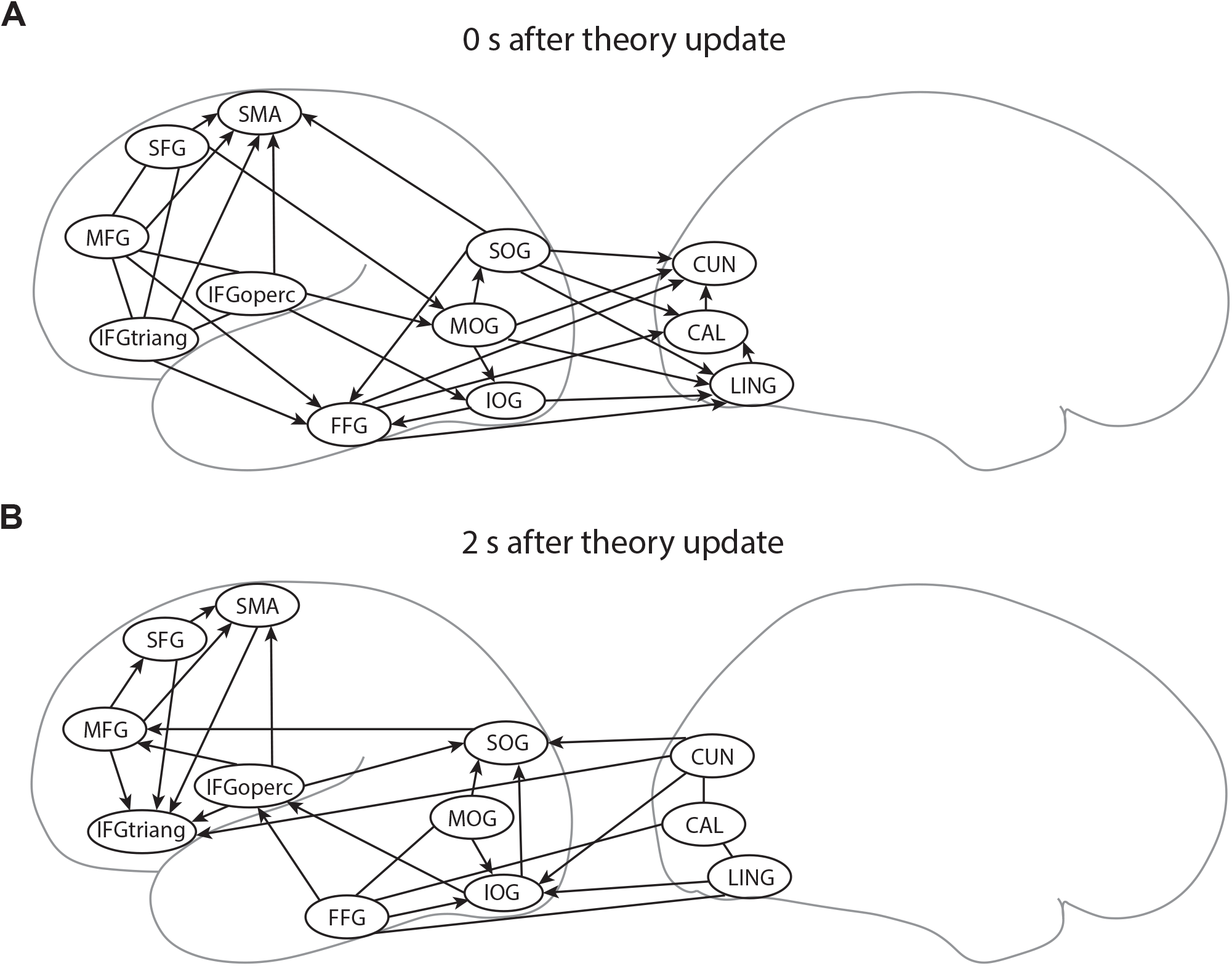
Effective connectivity during theory updating is consistent with predictive coding. A. Best-fitting effective connectivity pattern based on neural responses to theory update events estimated using beta series GLM. ROIs noted as in Figure 3B. B. Same results using neural responses 2 s after theory update events

## 8. Discussion

A longstanding question in neuroscience is how the brain represents the structure of the environment in order to support efficient learning and flexible generalization. One possible answer from cognitive science is that the brain learns a rich, abstract, causal model grounded in core cognitive concepts such as objects, relations, and goals, which is used to simulate the outcomes of different courses of action during planning (Lake et al., 2017; Tsividis et al., 2021; Pouncy et al., 2021; Pouncy and Gershman, 2022). We found support for this kind of theory-based RL using fMRI data from human participants learning to play different Atari-style games. The theory inferred by a theory-based RL agent can explain variance in inferior frontal gyrus and other prefrontal regions better than control models, suggesting that those regions encode theory-like representations above and beyond visual and model-free RL features. In an overlapping network of regions, including inferior frontal gyrus, occipital gyri, and fusiform gyrus, we found theory learning signals that could not be explained by visual events, motor actions, or theory-related nuisance variables, suggesting those regions play a role in theory inference. In a subset of those regions, we found evidence for separate learning signals for objects, relations, and goals, suggesting that the brain factorizes theory inference similarly to our theory-based RL agent. We additionally found that the striking overlap between theory coding and theory learning regions is not coincidental, with theory representations being activated following theory updates. This suggests that the theory is not stored as a persistent pattern of neural activity but is rather stored “silently” (Beukers et al., 2021), perhaps in the pattern of synaptic weights, and is only activated when updated by the theory inference circuitry. Finally, we found that the effective connectivity pattern during such updates is consistent with predictive coding (Rao and Ballard, 1999; Friston, 2005), with feedback connections conveying theory predictions and feedforward connections conveying theory prediction errors.

The idea that animals learn rich, structured the representations of their environments dates back to Tolman’s work on latent learning (Tolman and Honzik, 1930; Tolman, 1948). Tolman observed that rats were able to quickly find newly placed rewards in a maze after repeated unwarded exposures to the maze, leading him to hypothesize that this flexible generalization is supported by “cognitive maps” (Kaplan et al., 2017; Behrens et al., 2018; Boorman et al., 2021)—internal models of the world which allow animals to mentally search through space and find efficient paths to goals. Neural evidence for cognitive maps was famously identified in the hippocampus (O’keefe and Nadel, 1978), where place cells appear to encode an animal’s location in space. Subsequent studies found evidence that cognitive maps can represent nonspatial domains (Schuck et al., 2016; Constantinescu et al., 2016) and also appear in other parts of the brain (Walton et al., 2010; Rudebeck and Murray, 2011; Jocham et al., 2016) such as ventral prefrontal cortex which includes inferior frontal gyrus, a region our study implicates in theory coding. Our study did not find evidence for theory coding in the hippocampus, likely due to the fact at the theory on its own does not constitute a map *per se* but rather a set of abstract relational rules that, when grounded in a particular world state (such as a video game frame), can be used to predict future world states. We conjecture that the hippocampus and medial entorhinal cortex might be involved in such grounded representations, encoding a theory-based transition structure between concrete world states that directly supports planning, rather than the abstract theory itself.

Our findings resonate with recent studies that have used computational modeling to identify a more specific role for prefrontal cortex in representing and/or updating an internal causal model of the world (Donoso et al., 2014). In an fMRI study comparing model-based and model-free RL prediction errors, Gläscher et al. (2010) reported state prediction error signals—discrepancies between the observed state and the state predicted by the brain’s internal model, akin to theory prediction errors in our study – in similar prefrontal regions, particularly in bilateral inferior frontal gyrus. Another fMRI study by Lee et al. (2015) reported evidence of rapid, one-shot learning of causal associations driven by uncertainty encoded in ventrolateral prefrontal cortex, including inferior frontal gyrus. An fMRI study of causal structure learning from our lab (Tomov et al., 2018) found causal structure learning signals in a distributed bilateral network of regions, including inferior, middle, and superior frontal gyrus, regions in occipital cortex, and regions in the ventral stream such as fusiform gyrus. In that study, we also reported evidence of beliefs about causal structure being activated in response to feedback in a frontoparietal network of regions, including inferior frontal gyrus. Another study from our lab (Dorfman et al., 2021) reported evidence for beliefs about causal structure activated in inferior frontal gyrus during belief updating.

A separate line of work has implicated similar prefrontal regions in relational reasoning (Waltz et al., 1999; Krawczyk et al., 2011). Knowlton et al. (2012) unified some of these findings using a role-based relational reasoning model (LISA) according to which prefrontal cortex encodes abstract relational rules as distributed role-filler bindings at increasing levels of abstraction, from objects to relations to propositions, somewhat reminiscent of our HRR theory code. In LISA, rules are rapidly updated via spike-timing dependent plasticity in anterior prefrontal cortex and are activated in working memory by long-distance connections from semantic units in posterior cortex. This bears striking resemblance to our proposal, and suggests that theory-based reinforcement learning could serve as a unifying lens for results from the neuroscience literature on model-based reinforcement learning, causal inference, and relational reasoning. According to this view, these findings could be interpreted as signatures of the same theory inference machinery applied to different, narrower domains, with inferior frontal gyrus serving as the key locus of theory computation/storage in prefrontal cortex and posterior regions computing theory prediction errors for theory learning.

Video games have long served as microcosms in which to compare human and machine intelligence in naturalistic, complex environments (Mnih et al., 2015; Tsividis et al., 2017). Most closely related to our work is a recent study by Cross et al. (2021) in which fMRI data from human participants playing Atari games was analyzed using a deep RL network (DQN), a powerful model-free RL algorithm. The authors found evidence for DQN representations across a distributed network of regions, most notably in the dorsal visual stream and posterior parietal cortex. Despite similar methodology, there are crucial differences between our studies. The most critical difference is that we are interested in how people learn to play these games—an aspect of human behavior that is particularly well-captured by a theory-based RL model compared to model-free deep RL models—whereas Cross et al. (2021) are interested in the sensorimotor transformations that support gameplay after learning has plateaued. This in turn dictates important design decisions that differ between the two studies. Most importantly, we focus on games in which our prior work suggests that people seem to be model-based, and in particular that they seem to follow the predictions of theory-based RL, whereas Cross et al. (2021) focus on games in which people follow the predictions of the model-free DQN. As a result, our study includes more games played over shorter timescales with levels designed to maximize learning, less visually distinct features, and more complex rules. This could explain the relatively poor performance of our model-free RL control in matching human performance and brain activity.

However, our results are not mutually exclusive with those of Cross et al. (2021). Multiple studies have shown that brains employ a mix of model-free and model-based RL strategies (Gläscher et al., 2010; Daw et al., 2011; Lee et al., 2014; Kool et al., 2018). Indeed, the results from Cross et al. (2021) point to the dorsal stream, posterior parietal cortex, and motor areas as the loci of model-free sensorimotor transformations, whereas they report little evidence for model-free representations in prefrontal regions, and in particular they do not report any results in inferior frontal gyrus. In contrast, our results point to prefrontal cortex and inferior frontal gyrus in particular as the locus of theory encoding, and to occipital and ventral stream regions as the loci of theory learning computations; at the same time, we find little evidence for theory-based representations in the dorsal stream, posterior parietal cortex, or motor cortex. Thus the results from the two studies can be seen as complementary, pointing to hybrid architecture which includes both theory-based and model-free components. While EMPA in its current form is purely model-based, it can straightforwardly be extended to include learned policy and/or value components to help guide the search towards promising action plans. In the field of artificial intelligence, such hybrid approaches have recently achieved remarkable success in learning to play board games (Silver et al., 2017, 2018) and video games (Schrittwieser et al., 2020), suggesting that this could be a fruitful avenue for future neuroscience research.

Our effective connectivity analysis suggests that top-down information about the theory from prefrontal regions flows to occipital and ventral stream regions for predicting sensory inputs and that when a discrepancy occurs—a kind of theory prediction error—information flows the other way for updating the theory in prefrontal regions based on sensory input from occipital and ventral stream regions. This is broadly consistent with hierarchical predictive coding (Rao and Ballard, 1999; Friston, 2005): the idea that top-down (feedback) connections convey model predictions originating in higher cortical areas and shaping neural activity in lower cortical areas, which in turn compute prediction errors that are conveyed to higher areas via bottom-up feedforward connections for model updating. Despite this affinity, there are important differences between our proposal and traditional predictive coding accounts. First, the predictive coding interpretation only pertains to information flow between regions representing the learned theory and regions computing theory prediction errors. Importantly, it does not account for the processes of learning, planning, and exploration, which are core aspects of theory-based RL. Second, predictive coding models are usually employed in narrow domains, often focusing on simple problems of low-level perception (Rao and Ballard, 1999) or simple RL problems (Friston et al., 2013). In contrast, EMPA and theory-based RL more broadly focus on solving richer and more structured problems. Our approach considers perception and inference in the context of a complete modeling, planning and exploring agent; the models and plans generated by EMPA – and those generated by the brain – have more structure to them than those generated by standard predictive coding approaches. Finally, theory-based RL and predictive coding are frameworks at fundamentally different levels of description (Marr and Poggio, 1976): theory-based RL is a computational-level proposal of exploration, modeling, and planning based on Bayesian inference over intuitive theories (with EMPA being a particular algorithmic instantiation of it), whereas predictive coding is an implementation-level proposal of neural coding and dynamics of modeling and perception (Aitchison and Lengyel, 2017). Viewed in this light, our results suggest that the general predictive coding framework could be a promising starting point for studying theory predictions, theory prediction errors, and theory updating at the neural level. Future work could formally relate EMPA to particular predictive coding formulations, which could provide a richer theoretical framework for understanding the interplay between top-down and bottom-up inferential processes in the brain, as well as the interplay between model learning, exploration, and planning, relative to current predictive coding models.

One puzzling aspect of our results is the prevalence of visual regions, which raises the possible concern that our analysis was not selective enough to exclude visual confounds. This concern is partly addressed by our control analyses. In our encoding model analysis (Figure 3), we found that EMPA consistently outperformed all of our control models in prefrontal regions, but not in other cortical areas; indeed, in most other regions, EMPA was no better than PCA, suggesting that the theory effects in those areas could be partly explained by visual features. The theory update GLM (Figure 4) included visual nuisance regressors that showed a stronger effect in some regions, particularly in early visual areas, suggesting that those regions play a role in visual processing that is not specific to theory updating. Accordingly, we excluded early visual areas from reporting and follow-up analyses. Theory learning effects in higher visual areas could be partly explained by our effective connectivity results: according to the predictive coding interpretation, it is precisely visual regions that ought to compute theory prediction errors – discrepancies between theory-based predictions and sensory observations – which in turn serve as the basis for updating the theory in prefrontal regions. It is also worth noting that previous work on causal structure learning (Tomov et al., 2018) has also reported evidence for model updating in visual areas. Additionally, to some extent our experimental design already controls for visual confounds by having participants play the same level on repeat for 1 minute: if they do end up playing the same level for multiple episodes, most learning often occurs during the first episode(s), with the other episodes serving as implicit controls with nearly identical visual inputs but little-to-no theory learning. This idea could be taken further by having participants watch a replay of their own gameplay immediately after or in a subsequent scan session. We leave this kind of control study as future work.

Our study identifies key brain regions involved in theory encoding and theory updating, as well as the dynamic relationship between these processes. This can serve as a basis for further characterizing the neural architecture of theory-based RL. One important question pertains to theory coding: how exactly is the symbolic theory encoded in distributed patterns of neural firing and/or synaptic weights? Our study points to HRRs as one possibility, although we do not make strong theoretical commitments to HRRs, but merely use them is part of our encoding model. HRRs were originally proposed as a model of associative memory (Plate, 1995) and have since been used as a model of structured scenes in event memories (Franklin et al., 2020), making them a plausible candidate for a neural theory code. The theory coding question could be addressed by comparing encoding models using different theory codes.

Another open question pertains to theory learning: what algorithm does the brain employ to infer the theory and how is it implemented in neural circuits? Our study takes a step in addressing that question, indicating that theory inference might be factorized into separate object, relation, and goal updates, similarly to EMPA. However, unlike EMPA, which treats observations as deterministic and does not explicitly maintain probability distributions over theories, humans can account for the stochasticity of noisy observations and can maintain/update probabilities for multiple possible hypotheses (Donoso et al., 2014). An alternative to EMPA’s deterministic theory update is a particle-based approximation which explicitly maintains multiple hypothesized theories and their relative probabilities (Pouncy and Gershman, 2022). Both of these approaches maintain multiple hypotheses and progressively reduce uncertainty about the world model as the learner gets more data. Other possibilities come from deep model-based RL approaches (Schrittwieser et al., 2020), which can learn a model of the environment from scratch, or from deep meta-RL approaches (Duan et al., 2016; Wang et al., 2016), which use a model-free RL algorithm to learn a model-based RL algorithm. Future versions of EMPA could be extended with different inference algorithms to address this question.

A third open question pertains to the inductive biases which allow EMPA to learn as effectively as humans: where do they come from and how are they learned and represented by the brain? Work from our lab (Pouncy and Gershman, 2022) has proposed formalizing such inductive biases using a Markov logic network defining a prior distribution over theories. A follow-up study that explicitly manipulates participants’ biases could begin to address these questions.

In summary, our results are consistent with a neural architecture of theory-based RL in which theory representations in inferior frontal gyrus and other prefrontal regions are activated and updated in response to theory prediction errors computed in occipital and ventral stream regions, such as fusiform gyrus, in a way consistent with hierarchical predictive coding. Additionally, we hope that our work highlights the benefits of combining sophisticated, interpretable, end-to-end cognitive models such as EMPA with naturalistic experimental environments such as video games. By comparing the internal representations of such models with brain activity, researchers can begin to uncover how the brain learns and represents an internal model of the environment that supports adaptive behavior in complex, naturalistic tasks.

## 9. Acknowledgments

This research was supported by the Toyota Corporation, the Center for Brains, Minds and Machines (CBMM), funded by NSF STC award CCF1231216, and the Multi-University Research Initiative Grant (ONR/DoD N00014-17-1-2961). This work involved the use of instrumentation supported by the NIH Shared Instrumentation Grant Program award number S10OD020039. We acknowledge the University of Minnesota Center for Magnetic Resonance Research for use of the multiband-EPI pulse sequences. We are grateful to Alicia Chen, Yichen Li, Zhenglong Zhou, Daphne Cornelisse, Jiajia Zhao, Dimitar Karev, and Jason Ma for their help with data collection and initial prototyping.

## 10. Methods

### 10.1. Participants

Thirty-two healthy participants were recruited from the Cambridge, MA community: 15 female, 17 male, 19-36 years of age, mean age 24 ± 4 years, all right-handed and with normal or corrected-to-normal vision. The study was approved by the Harvard University Institutional Review Board and all participants gave informed consent. All participants were paid for their participation.

### 10.2. Experimental Design

Each participant played 6 different Atari-style games adapted from Tsividis et al. (2021) over the course of 6 scanner runs in a single session (Figure 2B). Six games were played across 6 scanner runs. Each run consisted of 3 blocks. Each block consisted of 3 levels of a given game. Each level was played on repeat for 1 minute total: if the episode ended before 1 minute had elapsed, a new episode began on the same level. Nine levels were played in total for each game. Scanner runs were grouped in 3 data partitions for cross-validation. Game order was pseudo-randomized such that each data partition contained one block of each game, ensuring that games and levels were balanced across partitions.

For each participant, games were randomly assigned names that were unrelated to the game rules (Archeplan, Deception Eagle, Dreams of Origins, Giants of Solitude, Questtide, Fuseville, Prime Origin). At the beginning of each block, the game name was shown for 2 s (Figure 2A). During an episode, the game name and the current score were displayed at the top of the screen. At the end of an episode, the outcome (“You WON!” or “You LOST!”) was shown at the bottom of the screen and the final frame was frozen for 2 s. Timing was adjusted such that each level was played for one minute total. After one minute, the current episode was interrupted with a “End of level” outcome (to distinguish it from a win or loss) shown for 2 s, unless the participant was already on a win/loss screen. There was a 10-s fixation cross at the beginning and end of each run to account for scanner stabilization and the hemodynamic lag, respectively. Each run was 566 s in total.

Following Tsividis et al. (2021), in order to avoid biasing learning with semantic priors based on object appearance, all games were played in “color-only” mode: all objects were visualized as colored squares with symbols on them. Objects of the same kind had the same color and symbol, while objects of different kinds had different colors and symbols. Color and symbol assignments were randomized across games and participants. The game descriptions were inspired by and/or drawn from the General Video Game AI (GVGAI) competition (Perez-Liebana et al., 2015) and expressed in the Video Game Description Language (VGDL; Schaul, 2013). Participants 1 through 11 played Chase, Helper, Bait, Zelda, Lemmings, Plaque Attack. Participants 12 through 32 played the same games, except for Plaque Attack which was replaced by Avoid George. Each game had 5 actions: move left, move up, move down, move right, action key. The levels were designed to ensure continuous learning about the game rules. Specifically, different levels involved different object layouts and later levels involved game rules that earlier levels did not. All game and level descriptions are available at https://github.com/tomov/RC_RL/tree/fmri/fmri_all_games.

Participants were told that they would be playing a sequence of Atari-style games with different rules and that they will have to learn the rules of each game from experience. The game and level order and timing was explained to them (Figure 2B), as well as that they would be playing all games in color-only mode and what that is. Specifically, they were told that the colors, symbols, and game names convey no information about the game rules, except that objects of the same kind look the same in a given game. They were also told that colors and symbols in one game convey no information about objects in another game. All participants were paid a base of $80 for their participation. Additionally, to incentivize learning, we paid participants a bonus based on performance. Specifically, for each participant, we randomly chose a level and paid them the maximum score they achieved (in dollars) at the end of any episode on that level, counting only episodes which they won. If they never won that level, the bonus was $0. This bonus scheme was explained to them in detail. They were also told that it is meant to encourage efficient learning and gameplay: they should aim to maximize the score and win each level within 1 minute.

In the scanner, participants played using a 5-finger button box, with each button corresponding to a game action (index finger = move left, middle finger = move up, ring finger = move down, pinky finger = move right, thumb = action key). Before entering the scanner, participants practiced by playing 3 levels (1 block) of a different game (Sokoban) on the laptop using a similar key setup. This game was not played in the scanner. Overall, the entire scan session took 2.5 hrs per participant, 1.5 hrs of which was spent in the scanner, 1 hr of which was spent on BOLD acquisition and gameplay.

### 10.3. fMRI Data Acquisition

We followed a similar protocol to our previous work (Tomov et al., 2020). All participants were scanned using a 3T Siemens Magnetom Prisma MRI scanner with the vendor 32-channel head coil (Siemens Healthcare, Erlangen, Germany) at the Harvard University Center for Brain Science Neuroimaging. A T1-weighted high-resolution multi-echo magnetization-prepared rapid-acquisition gradient echo (ME-MPRAGE) anatomical scan (Van der Kouwe et al., 2008) of the whole brain was acquired for each participant prior to any functional scanning: 176 sagittal slices, voxel size = 1.0 × 1.0 × 1.0 mm, TR = 2530 ms, TE = 1.69–7.27 ms, TI = 1100 ms, flip angle = 7^°^, FOV = 256 mm. Functional images were acquired using a T2*-weighted echo-planar imaging (EPI) pulse sequence that employed multiband RF pulses and Simultaneous Multi-Slice (SMS) acquisition (Moeller et al., 2010; Feinberg et al., 2010; Xu et al., 2013). We collected 6 functional runs for each participant, each with 283 timepoints (Figure 2B). Scan parameters: 87 interleaved axial-oblique slices per whole-brain volume, voxel size = 1.7 × 1.7 × 1.7 mm, TR = 2000 ms, TE = 30 ms, flip angle = 80°, in-plane acceleration (GRAPPA) factor = 2, multi-band acceleration factor = 3, FOV = 211 mm. Functional slices were oriented to a 25° tilt towards coronal from AC-PC alignment. The SMS-EPI acquisitions used the CMRR-MB pulse sequence from the University of Minnesota.

All 32 participants were included in the analysis. Scanner runs with excessive motion (> 3 mm translation or > 3° rotation) were excluded from the analysis.

### 10.4. fMRI Preprocessing

Following our previous work (Tomov et al., 2020), we preprocessed functional images using the `SPM12 MATLAB` toolbox (Wellcome Department of Imaging Neuroscience, London, UK). Each functional scan was realigned to correct for small movements between scans, producing an aligned set of images and a mean image for each participant. The high-resolution T1-weighted ME-MPRAGE images were then co-registered to the mean realigned images and the gray matter was segmented out and normalized to the gray matter of a standard Montreal Neurological Institute (MNI) reference brain. The functional images were then normalized to the MNI template (resampled voxel size 2 mm isotropic), spatially smoothed with a 8-mm full-width at half-maximum (FWHM) Gaussian kernel, high-pass filtered at 1/ 128 Hz, and corrected for temporal autocorrelations using a first-order autoregressive model.

### 10.5. EMPA

A detailed technical description of EMPA can be found in Tsividis et al. (2021), which we summarize here. EMPA learns a model (or theory), *θ*, of each game expressed in VGDL (Schaul, 2013). VGDL breaks down the game rules into three different components corresponding to core aspects of human intuitive theories (Lake et al., 2017; Carey, 2000): objects (sprites), relations (interactions) between objects, and goals.

A VGDL game description consists of a SpriteSet, *θ*_*S*_, which specifies the type, appearance, and dynamic properties each object (e.g., “red objects chase the avatar at a speed of 3 squares per second”); an InteractionSet, *θ*_*I*_, which specifies what happens when two objects interact (e.g., “when a red object collides with the avatar, the avatar dies”); and a TerminationSet, *θ*_*T*_, which specifies the win/loss conditions of the game (e.g., “when the avatar dies, the game is lost with a score of 0”). A VGDL description thus procedurally defines a Markov Decision Process: the state at every timestep is described by the object instances and locations, the avatar’s internal state, and any events due to collisions between pairs of objects; the transition function is defined by the SpriteSet, the InteractionSet, and the TerminationSet; and the reward function is defined by the InteractionSet and the TerminationSet.

EMPA learns the rules of each game by inferring a probability distribution over the space of possible VGDL theories, Θ, from experience using Bayesian inference:

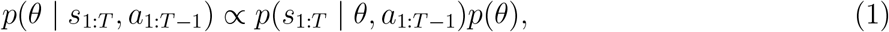

where *θ* = (*θ*_*S*_, *θ*_*I*_, *θ*_*T*_) is the inferred theory describing the game rules, *T* is the current timestep, *s*_1:*T*_ is the history of observed states, *a*_1:*T* −1_ is the history of avatar actions, and *p*(*θ*) is a minimum description length prior favoring simpler theories.

To choose actions, EMPA uses the maximum *a posteriori* theory, *θ*^*^, together with a simulation-based planner that searches for action sequences that lead to rewarding outcomes under *θ*^*^. Specifically, EMPA pursues exploitative goals that lead to winning (according to 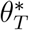), as well as exploratory goals that reduce the uncertainty in *θ* (e.g., inducing an unobserved collision). Pursuit of these sparse goals is aided by subgoals, which represent partial progress towards goals (e.g., “3 blue objects remaining”), and goal gradients, which represent preferences for states that are spatially closer to achieving a subgoal (e.g., “the closest blue object is 3 squares away”). Planning is further aided by state pruning and re-planning based on prediction errors, as described in Tsividis et al. (2021).

The code for EMPA will be available at https://github.com/tsividis/vgdl upon publication.

### 10.6. DDQN

Following Tsividis et al. (2021), as a control model we trained a deep reinforcement learning network (DDQN) based on the public repository https://github.com/dxyang/DQN_pytorch with parameter settings *α* = 0.00025, *γ* = 0.999, *τ* = 100, experience-replay max = 50, 000, batch size = 32, and image input recrop size = 64 × 64 × 3. The exploration parameter, ϵ, was annealed linearly from 1 to 0.1 using a decay rate of 200 steps. Following (Mnih et al., 2015), the DDQN had 3 convolutional layers (`conv1`: 32 filters with size = 8 × 8 and stride = 4; `conv2`: 64 filters with size = 4 × 4 and stride = 2; conv3: 64 filters with size = 3 × 3 and stride = 1), followed by a fully connected layer (`linear1`: 512 units), followed by the output layer (`linear2`: 5 units). Each convolutional layer was followed by batch normalization and linear rectification (ReLU). ReLU units also followed the fully connected layer. The input was a 64 × 64 × 3 scaled game frame with 3 color channels (RGB). To ensure a fair comparison with EMPA, we pretrained a separate DDQN for each game using a VGDL environment for 100 epochs of 250, 000 steps. Levels were alternated across epochs to ensure exposure to all levels. Specifically, in each epoch, the DDQN was trained on a given level for one or more episodes, restarting the level if it was won or lost. During epoch 1, we trained on level 1, during epoch 2, we trained on level 2, and so on, starting over from level 1 after level 9. We used the same pretrained DDQNs for both the behavioral and the neural analyses.

The code for the DDQN is available at https://github.com/tomov/RC_RL.

### 10.7. Generative play

To compare human performance with EMPA and DDQN performance, we valuated the models on the same games and levels as the human participants. We simulated each participant with EMPA by having a separate EMPA instance play all levels of each game generatively, in order. As with human participants, each level was played for 1200 frames (60 sec at 20 Hz), restarting the level if won or lost before that. Similarly to humans, performance was evaluated based on the expected bonus payout, namely the maximum per-level winning score, averaged across all levels and games. We simulated 32 participants independently, each simulation corresponding to a single human participant. We similarly simulated 32 participants with the pretrained DDQNs. Note that, unlike the DDQNs, EMPA does not require pretraining.

### 10.8. Encoding model analysis

To compare EMPA theories to brain activations, we used an encoding model (Naselaris et al., 2011; Güçlü and van Gerven, 2015; Cross et al., 2021) that maps EMPA theory embeddings to BOLD signal (Figure 3A). For each participant, we first replayed the sequence of states, actions, and rewards from their gameplay in the scanner through EMPA, using a separate EMPA instance for each game. This produced an EMPA theory for each frame, corresponding to the theory that EMPA would have inferred at that timepoint if it had observed the same sequence of events as the participant. We embedded each theory in a vector space using holographic reduced representations (HRRs; see below), resulting in a sequence of HRR embeddings. To account for the stochasticity inherent in HRRs, we independently generated 100 such sequences, each with a different random HRR initialization. Each sequence was convolved with the canonical hemodynamic response function from `SPM` and subsampled at the scanner frequency (TR = 2 s, or 0.5 Hz).

For each voxel, we predicted the BOLD signal with Gaussian process (GP) regression (see below) using cross-validation across the 3 data partitions (Figure 2B). We quantified accuracy by correlating the predicted with the actual BOLD signal for each partition, averaging the resulting Pearson correlation coefficients across partitions, and Fisher z-transforming the result to obtain a single predictivity score *z* for that voxel. To aggregate across participants, we performed a two-sided t-test against 0 across participants for each voxel, producing a group-level statistical map (t-map). Following our previous work (Tomov et al., 2020), we thresholded single voxels at *p* < 0.001 and applied cluster family-wise error (FWE) correction at significance level *α* = 0.05. We visualized the corrected t-maps using the `bspmview` toolbox in `MATLAB`.

Anatomical regions of interest (ROIs) were extracted by cross-referencing the peak voxels in each cluster (up to 3 peaks peaks per cluster, minimum 20 voxels apart) with the automated anatomical labeling atlas (AAL3 atlas; Rolls et al., 2020). Confirmatory ROI analyses were performed using bilateral anatomical ROIs from all models (see Control models below). In a given ROI, for each participant we computed the fraction of significant voxels as the number of voxels with a significant Pearson correlation at the *α* = 0.05 significance level, divided by the total number of voxels in the ROI. We compared models in each ROI using Wilcoxon signed rank tests across participants. To aggregate ROIs into ROI groups (Figure S3C,D), we simply merged ROIs from a given cortical region into a single “macro-ROI” and performed the same analysis.

We similarly applied GP regression with our control models.

### 10.9. Gaussian Process regression

For the encoding model we used Gaussian process (GP) regression (Williams and Rasmussen, 2006), a nonparametric method for predicting values of unseen data points based on similarity with observed data points. Ridge regression – a more commonly used encoding model (Driscoll et al., 2017; Cross et al., 2021) – can be derived as a special case of GP regression. However, unlike ridge regression, GP regression avoids the need to fit weights to individual HRR components (which by design are random) and allows for straightforwardly accounting for the randomness of HRRs.

To justify the use of GP regression, first consider the standard general linear model (GLM) formulation:

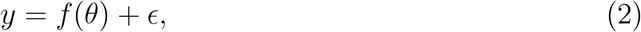

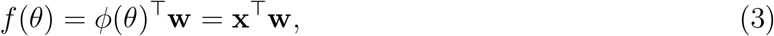

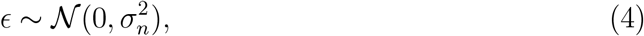

where *y* is the neural signal at a given time point, *θ* is the EMPA theory, *ϕ*(*θ*) = **x** is the HRR embedding of *θ*, **w** are the component weights (often referred to as beta coefficients), and *ϵ* is Gaussian noise with zero mean and variance 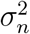. Such GLMs are routinely used to fit brain data and the resulting weights **w** – often fit using maximum likelihood estimation – are used to interpret whether a given feature is represented in brain activity.

High-dimensional feature spaces pose a challenge to this approach, as the weights might be underconstrained. One way around this is to impose a prior distribution on the weights:

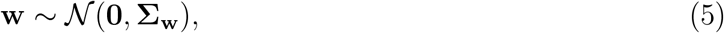

where **Σ**_*w*_ is the prior weight covariance matrix. The maximum *a posteriori* solution to this Bayesian linear regression problem is equivalent to ridge regression, where a regularization term that constrains the weights arises naturally from the weight prior.

The challenge with applying ridge regression is that HRR embeddings are random, which 1) renders the weights meaningless, and 2) necessitates averaging over that randomness. These issues can both be addressed by GP regression. First, the predicted neural signal *ŷ*_*_ for a theory *θ*_*_ can be directly computed in closed form from the training data *θ*, **y** (Williams and Rasmussen, 2006), bypassing the need to compute the weights:

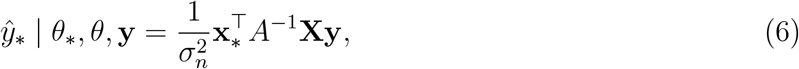

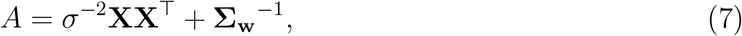

where *θ* = [*θ*_1_, *θ*_2_, *θ*_3_, …]^⊤^ and **y** = [y_1_, y_2_, y_3_, …]^⊤^ are the training theory and neural activation sequences, respectively, *θ*_*_ and *y*_*_ are the held-out theory and neural activation, respectively, **x**_*_ = *ϕ*(*θ*_*_) is the HRR embedding of the held-out theory, and **X** = [**x**_1_, **x**_2_, **x**_3_, …] = [*ϕ*(*θ*_1_), *ϕ*(*θ*_2_), *ϕ*(*θ*_3_), …] is the training data design matrix. Note that we are only using the posterior means and omitting the variances for ease of exposition.

This can be further rearranged by applying the “kernel trick” (Williams and Rasmussen, 2006), resulting in GP regression:

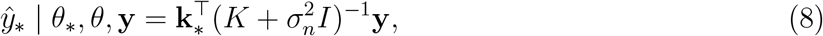

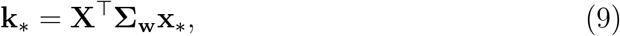

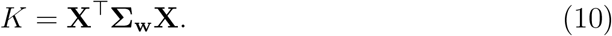

Here, the covariance matrix (or kernel) *K* quantifies the similarity between every pair of theories in the training data, and the covariance vector **k**_*_ quantifies the similarity between every training theory and the held-out theory. In our case, they were computed based on the HRR design matrix **X**, but in principle we could use a similarity metric that does not rely on explicitly computed features. We used **Σ**_**w**_ = *I*, so our similarity metric for each pair of theories was effectively the dot product of their HRR embeddings. Intuitively, equation 8 says that the predicted held-out neural activation is the average of the training neural activations, weighted by the similarity between the corresponding training theories and the held-out theory.

Finally, we can account for the randomness of HRRs by marginalizing over different HRR embedding functions *ϕ* resulting from different HRR initializations:

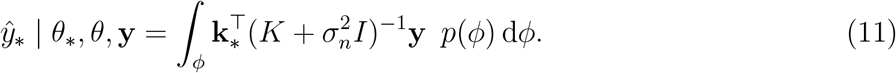

From the central limit theorem and the stochasticity of HRRs, the resulting distributions of *K* and **k**_*_ are approximately Gaussian, so we chose to simplify further by approximating them using Dirac delta functions around their means, 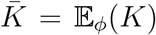 and 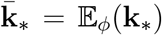, yielding the final GP formulation that we used:

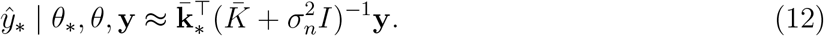

We used a sampling approximation for 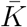 and 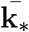 by averaging over the kernels for 100 different HRR initializations. In practice, during cross-validation, we had a set of held-out data points *θ*_*_, **y**_*_ (rather than a single data point) with a corresponding covariance matrix *K*_*_ between the training and held-out data points. So for each HRR initialization we computed a single kernel for all three data partitions, averaged the kernels across different HRR initializations, and then selected submatrices of the average kernel to get 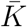 fold. and 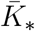 accordingly for each cross-validation

Our initial results indicated that our analysis is confounded by game identity: it produces nearly identical results to those of a simple model were the feature vector **x** is a 6-dimensional one-hot vector representing the game currently being played (Figure S2). To address this, we projected out game identity from the BOLD signal and the model:

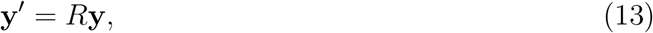

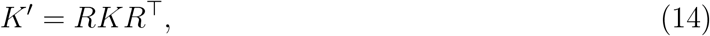

where 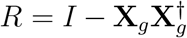 is the residual forming matrix for the game identity encoding model defined by the design matrix **X**_*g*_ († denotes the Moore-Penrose pseudoinverse). This is equivalent to using the residuals from a game identity GLM fit to the BOLD signal.

### 10.10. Holographic reduced representations

We embedded EMPA theories in a vector space using holographic reduced representations (HRRs; Plate, 1995), a kind of vector symbolic architecture (Gayler, 2004) that can represent compositional structure in distributed form. HRRs were originally proposed as a model of associative memory and have since been used for modeling structured memories of past events (Franklin et al., 2020). HRRs use circular convolution to associate pairs of items, represented by vectors, and addition to create bags of associations. The resulting vectors can be further associated or grouped together to represent higher-order compositions. Individual items could be extracted from the resulting vector using circular correlation, although we do not take advantage of this in our work.

An EMPA theory consists of three components – objects (SpriteSet), relations (InteractionSet), and goals (TerminationSet) – that we embed separately and then combine into a single vector (Figure S1).

The SpriteSet is a set of object (sprite) classes, each consisting of a set of properties with given values (e.g., `type=Missile, color=blue, speed=slow`). The base vectors corresponding to properties (e.g., type) and their values (e.g., `Missile`) are drawn from isotropic D-dimensional Gaussian distributions 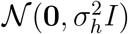, where 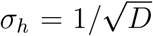, bound together using circular convolution, and added to produce the vector for the corresponding sprite class (e.g., `c3 = type` ⊛ `Missile` + `color` ⊛ `blue` + `speed` ⊛ `slow`). The vectors for different sprite classes are scaled to unit length and added together to produce the SpriteSet vector, which is also normalized to unit length.

The InteractionSet is a set of relations (interactions) between pairs of sprite classes, each describing the outcome of an interaction. Each interaction has three key properties: an agent sprite class, a patient sprite class, and an interaction type describing the outcome of the interaction (e.g., `killObject`). In addition, there may be other optional properties (e.g., `scoreIncrement`). As with the SpriteSet, the base vectors for properties and their values are drawn from D-dimensional isotropic Gaussian distributions, with the exception of values for agent and patient vectors which are the SpriteSet vectors for the corresponding sprite classes. The property and value vectors are bound together and added to produce the interaction vector (e.g., `i3 = patient` ⊛ `c0` + `agent` ⊛ `c3` + `interaction` ⊛ `killObject`). The vectors for different interactions are normalized to unit length and added together to produce the InteractionSet vector, which is also normalized to unit length.

The TerminationSet is a set of exploitative goals (termination conditions) and exploratory goals. Each termination condition has a type (e.g., `counter`), a sprite class, an outcome (e.g., `loss`), as well as any additional properties (e.g., `count`). Exploratory goals have two sprite classes whose interaction is yet unobserved, as well as other optional properties. As with the InteractionSet, the base vectors for properties and their values are drawn from D-dimensional isotropic Gaussian distributions, with the exception of spite classes whose vectors are the corresponding SpriteSet vectors. The property and value vectors are bound together and added to produce the goal vector (e.g., `t0` = `type` ⊛ `counter` + `sprite` ⊛ `c0` + `outcome` ⊛ `loss`). The values for different goals are normalized to unit length and added together to produce the TerminationSet vector, which is also normalized to unit length.

The resulting SpriteSet, InteractionSet, and TerminationSet vectors are finally added to produce the theory vector, which is also normalized to unit length. Following Plate (1995), we chose the dimension D of the vectors as:

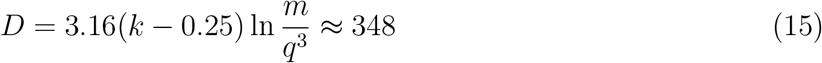

Where *k* = 10 is the number of stored vectors, *m* = 10 is the vocabulary size, and *q* = 0.05 is the probability of retrieval error.

Note that we are not making a strong commitment to HRRs as a neural code. Specifically, we are not testing the hypothesis that the brain encodes theories in a form similar to HRRs; rather, we are using HRRs as a cognitively plausible theory embedding to construct the theory similarity kernel *K* for our encoding model. We leave the question of theory coding as the topic of future work.

### 10.11. Control models

We performed a similar encoding model analysis with 3 control models:

- DDQN, to account for model-free RL representations,
- PCA, to account for low-level visual representations,
- VAE, to account for higher-level state representations.

Deep RL networks (DQNs) have achieved human-level performance on Atari games (Mnih et al., 2015) and have been put forward as an account of human model-free RL in complex domains (Cross et al., 2021). Following Tsividis et al. (2021), we used a double DQN (DDQN), which is a version of the original DDQN with improved convergence properties (van Hasselt et al., 2016). We ran the sequence of frames (scaled to 64 × 64 × 3), actions, and rewards from each participant through DDQNs pre-trained for the corresponding games, as described above. For each frame, we concatenated the activations from all layers into a single feature vector. The resulting feature vector sequences were fed through the same analysis pipeline as the EMPA theory embeddings (Figure 3A).

Principal component analysis (PCA) has been used to explain brain activity in the visual pathway (Olshausen and Field, 1996; Chang and Tsao, 2017) and has been utilized as a control model for human RL in Atari games (Cross et al., 2021). We first extracted principal components from 430,000 randomly chosen frames (scaled to 64 × 64 × 3) across all participants. We used the incremental PCA algorithm from the `sklearn Python` library with a batch size of 10,000. We then projected the frame sequence from each participant’s gameplay on to the top 100 principal components and fed the resulting feature vectors through the same analysis pipeline as the EMPA theory embeddings.

Variational autoencoders (VAEs) extract a latent representation of an input space by learning to compress and then reconstruct the input data using a deep neural network (Mohamed and Jimenez Rezende, 2015; Watter et al., 2015; Higgins et al., 2017). VAEs have also been used as a control model for human RL in Atari games (Cross et al., 2021). We used an open-source VAE implementation (https://medium.com/dataseries/variational-autoencoder-with-pytorch-2d359cbf027 The encoder had 3 convolutional layers (`conv1` : 8 filters with size = 3 × 3 and stride = 2; `conv2` : 16 filters with size = 3 × 3 and stride = 2; `conv3`: 32 filters with size = 3 × 3 and stride = 2), followed by 2 fully connected layers (`linear1` : 128 units, `linear2` : 128), followed by the bottleneck layer (latent: 128 units). The decoder had a correspondingly inverted architecture, with 2 fully connected layers followed by 3 convolution transpose layers. The VAE was trained by maximizing the evidence lower bound (ELBO) on the marginal likelihood of the training data. As with PCA, we trained on 430,000 random frames across all participants. We used batch size = 256 and trained for 1,000 epochs using the Adam optimizer with learning rate = 0.001 and weight decay = 10^−5^. We then ran the frame sequence from each participant’s gameplay through the VAE and used the bottleneck activations as the feature vectors which were fed through the same analysis pipeline as the EMPA theory embeddings.

### 10.12. GLM analyses

To look for brain regions sensitive to theory updates, we employed a standard GLM approach using SPM12 (Figure 4A). We created a GLM with impulse regressors at time points when the theory inferred from EMPA changed (*θ*_*t*_ ≠ *θ*_*t*−1_). We also included nuisance regressors for visual and motor confounds, variables relevant for theory updating, as well as motion estimates derived from realignment and run-specific intercepts (Table S3). All regressors were convolved with the canonical hemodynamic response function. As in our previous work (Tomov et al., 2020), group-level statistical maps were thresholded at p < 0.001 and cluster FWE corrected at *α* = 0.05.

As with the encoding model, ROIs were extracted by cross-referencing the peak voxels from the group-level t-map (up to 3 peaks peaks per cluster, minimum 20 voxels apart) with the AAL3 atlas (Rolls et al., 2020). For our confirmatory analysis, we used all anatomical ROIs with an average beta coefficient for theory updating which was significantly different from zero across participants (Figure S4). We generated PETHs for a given participant and ROI by taking the 20-s (10 TRs) BOLD timecourse following every theory update event, averaged across all voxels in the ROI, and subtracting a baseline BOLD signal averaged over the preceding 4 s (2 TRs) to obtain the change in BOLD signal in response to theory updating. The resulting traces were averaged across theory update events and aggregated across participants to obtain the final PETHs (Figure 4C). The same analysis was performed for the control events. To directly compare the change in BOLD signal in response to different kinds of events (Figure 4D), we averaged the BOLD timecourse within the 20-s window following each event before averaging across event instances and aggregating across participants.

### 10.13. GLM comparison

To identify regions which are sensitive to different update events, we constructed 4 additional GLMs analogous to the theory update GLM: 3 GLMs for single component updates (objects, relations, and goals, respectively) and a single GLM with separate regressors for all three component updates (Figure 5A). Following our previous work (Tomov et al., 2018), we compared GLMs using random effects Bayesian model selection (Rigoux et al., 2014). We approximated the log model evidence as LME = -0.5 * BIC, where BIC is the Bayesian information criterion based on the maximum likelihood estimate of the GLM parameters. This quantifies how well the GLM fits the BOLD signal in the ROI for a given participant (penalizing for model complexity), with lower values indicating a better fit. We report the protected exceedance probability (PXP), which is the posterior probability that a given model is most prevalent in the population (Table 1).

### 10.14. Theory activation timecourse

We generated the overlap between our theory representation and theory updating t-maps by taking the voxels that were significant in both group-level t-maps (Figure 6A). To generate PETHs with predictivity scores (Figure 6B) for a given ROI and participant, we first obtained a predictivity timecourse by computing the Fisher z-transformed Pearson correlation between the predicted and actual pattern of BOLD activity across voxels at each TR. We then proceeded in a similar fashion to the BOLD PETHs described above: the 20-s predictivity timecourses following theory updates were baseline-subtracted (average of preceding 4 s), averaged across theory update events, and aggregated across participants to obtain the PETHs. The same analysis was performed for the control events. As with the BOLD PETHs, to directly compare the change in predictivity score in response to different kinds of events (Figure 6C), we averaged the predictivity timecourse within the 20-s window following each event before averaging across event instances and aggregating across participants. When performing this analysis for separate theory component updates (Figure S6), we used predictivity scores from encoding models fit separately for objects, relations, and goals, respectively.

### 10.15. Effective connectivity

Following our previous work (Dorfman et al., 2021), we investigated the pattern of effective connectivity between brain regions during theory updating using structural equation modeling (Spirtes, 2005; Ramsey et al., 2010; Poldrack et al., 2011). We constructed a beta series GLM with separate impulse regressors for individual theory update events. Since the BOLD signal is highly autocorrelated, which violates the structural equation modeling assumptions, we only included events that are at least 10 s apart, using a rolling window starting from the first theory update event in each run. The resulting beta coefficients are estimates of the instantaneous neural activity at each theory update event. For each ROI, we averaged the estimates across voxels. We searched the space of connectivity patterns using the IMaGES (independent multiple-sample greedy equivalence search) algorithm (Ramsey et al., 2010; Poldrack et al., 2011) from the TETRAD software package for causal modeling (Scheines et al., 1998). IMaGES is a version of greedy equivalence search (GES; Meek, 1997), which starts with an empty causal graph and greedily adds edges that improve the fit to the data according to the BIC. IMaGES extends GES to multiple datasets (e.g., multiple fMRI participants) by averaging the BICs across datasets. To find the effective connectivity pattern 2 s after theory updating, we performed the same analysis except with all theory updates shifted back by 2 s.

## 11. Supplemental Information

**Figure S1:**
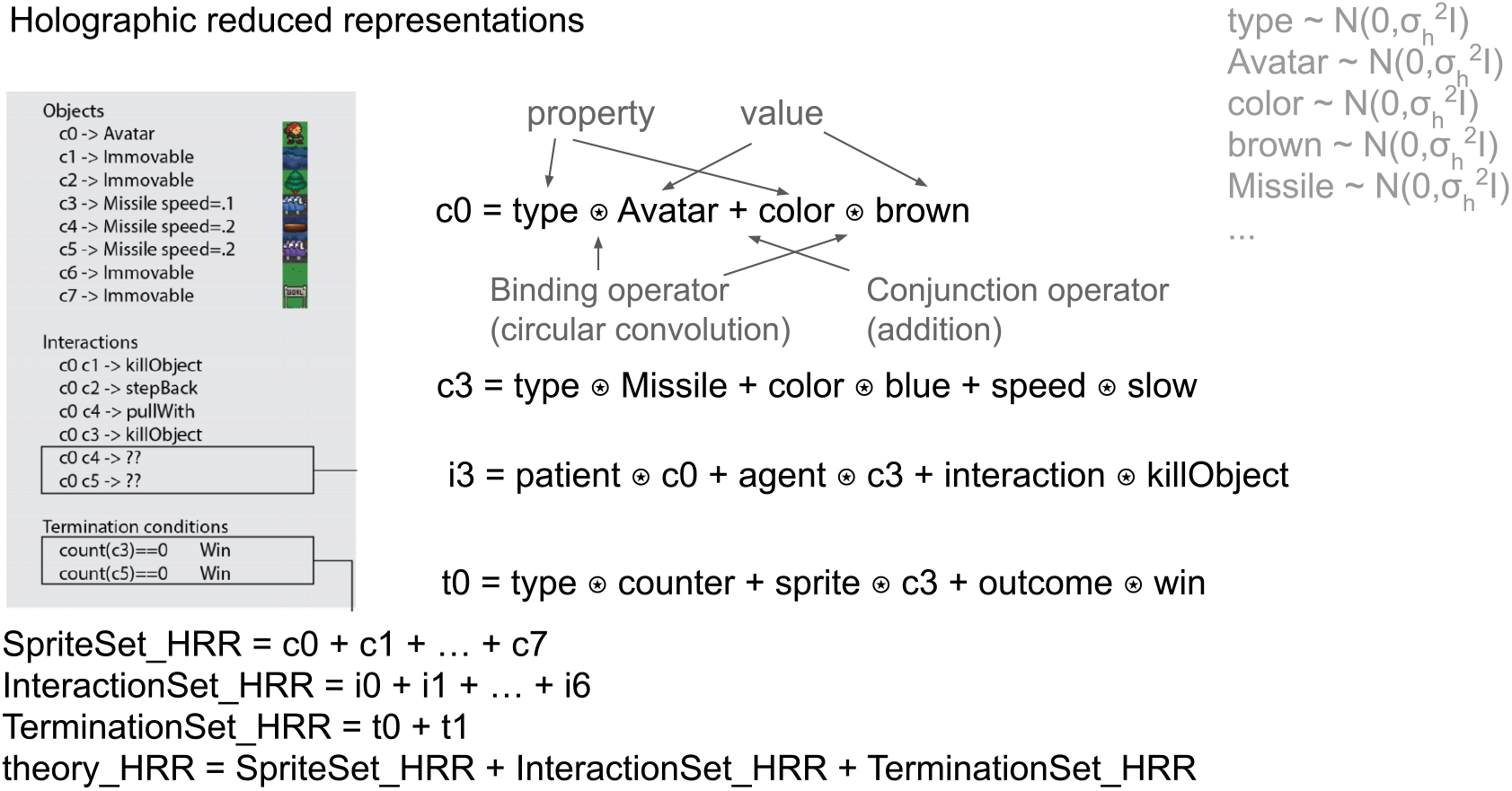
Illustration of HRRs. Related to Figure 3A.

**Figure S2:**
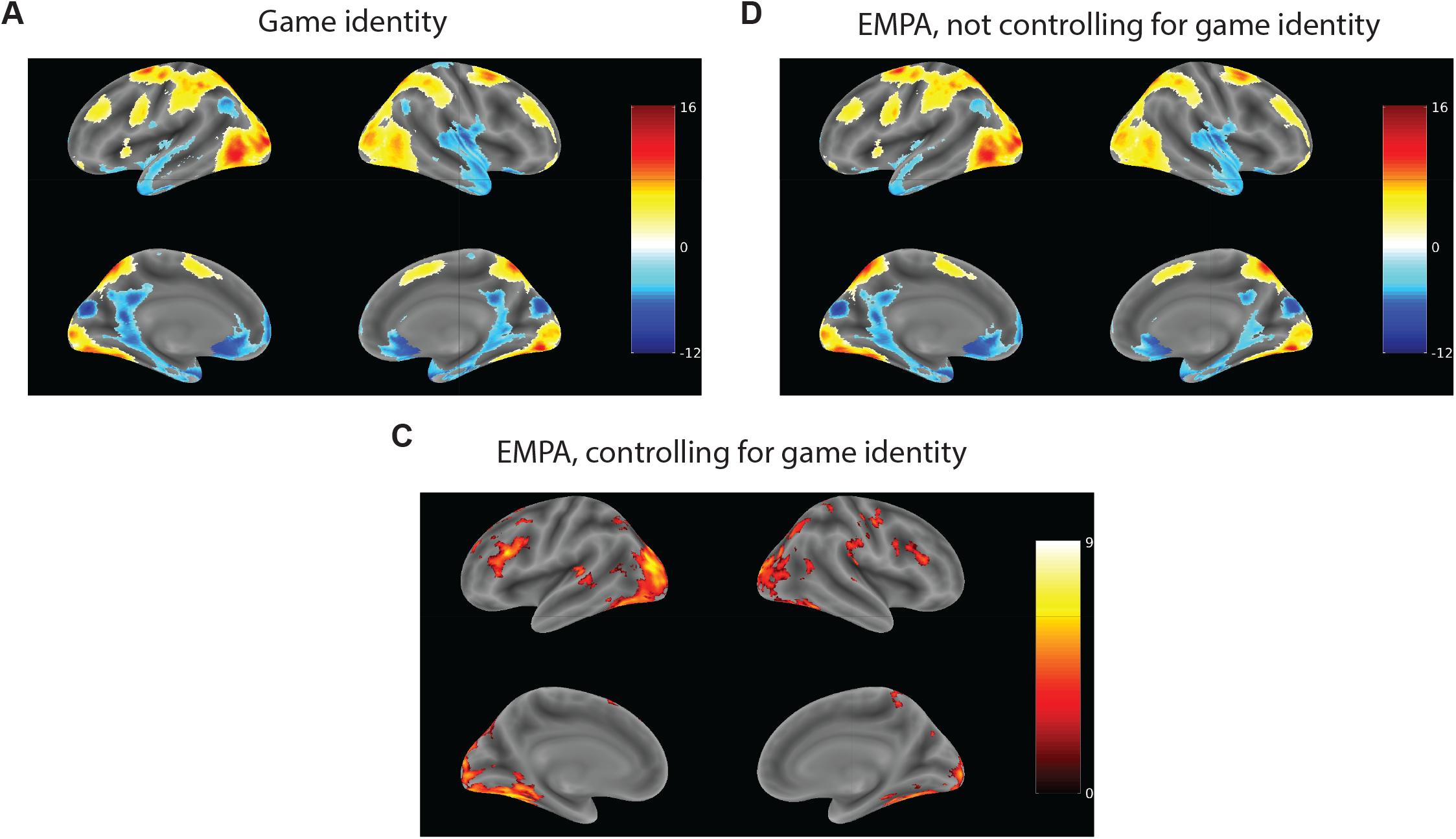
Controlling for game identity. Related to Figure 3B. A. Group-level t-map from encoding model analysis using one-hot game identity feature vectors. Notation as in Figure 3B. B. Group-level t-map from theory encoding model (Figure 3A) without controlling for game identity. C. Group-level t-map from theory encoding model (Figure 3A), controlling for game identity. Surface view corresponding to slices from Figure 3B.

**Figure S3:**
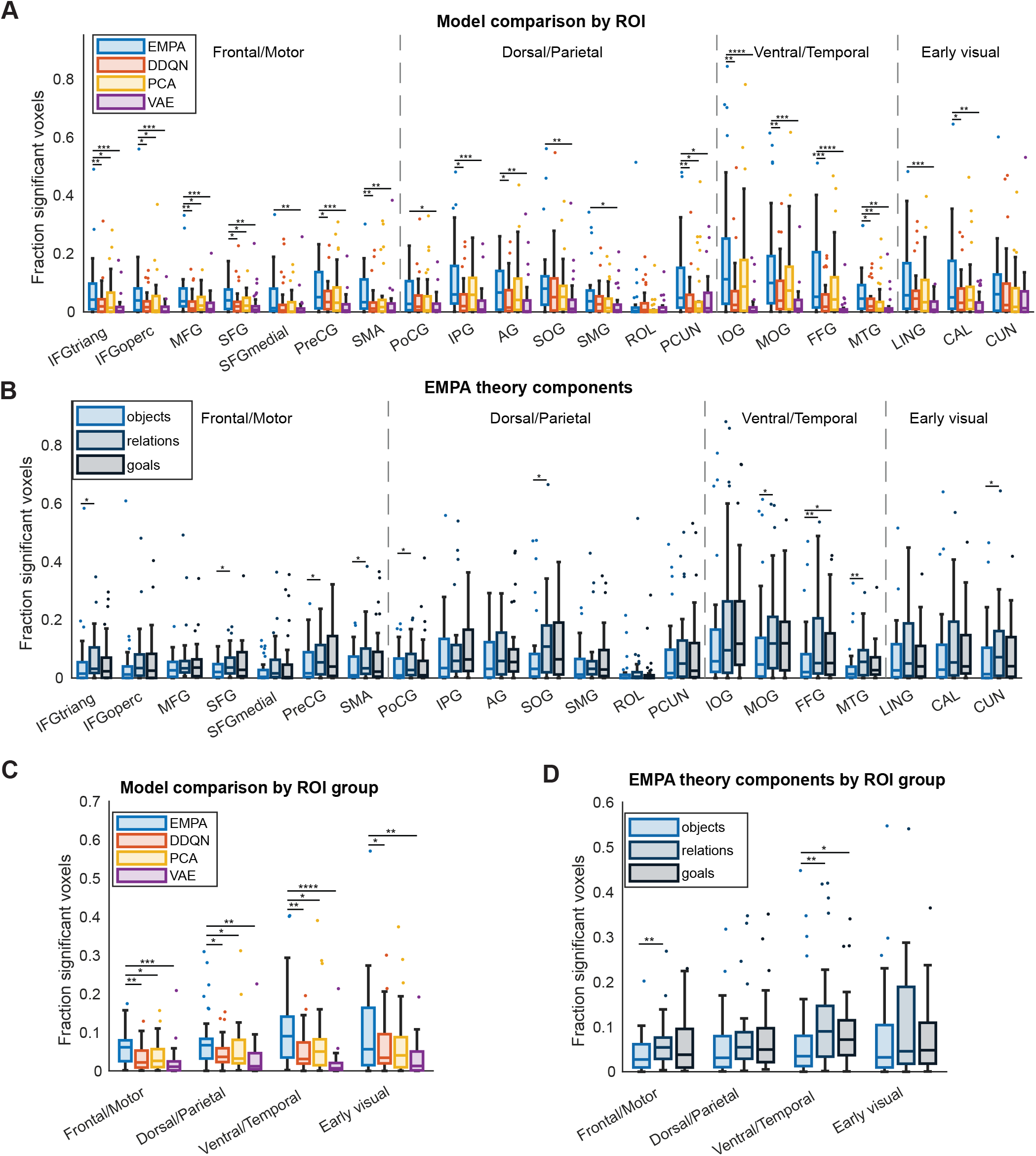
Encoding model results with outliers visualized. Related to Figure 3C. A. Fraction of voxels with significant correlation between predicted and actual BOLD for different models. Notation and results as in Figure 3C, except with outliers visualized as dots. B. Same as A, except for individual theory components. C. Results from A aggregated across larger cortical areas. D. Results from B aggregated across larger cortical areas.

**Figure S4:**
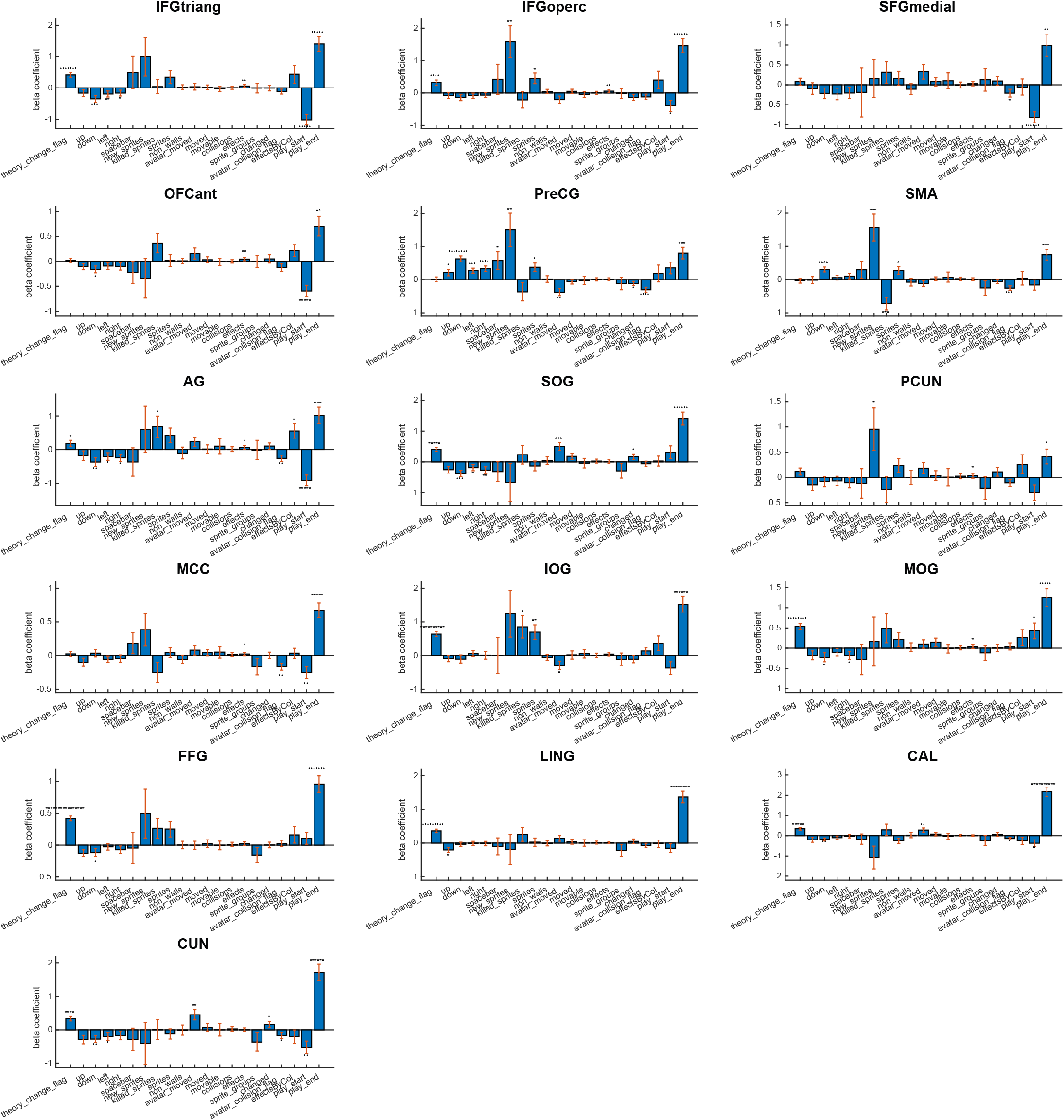
Beta coefficients from theory update GLM. Related to Figure 4. * - *p <* 0.05, ** - *p <* 0.01, *** - *p <* 0.001, **** - *p <* 0.0001, ***** - *p <* 0.00001, ****** - *p <* 0.000001, ******* - *p <* 0.0000001 (two-sided t-tests).

**Figure S5:**
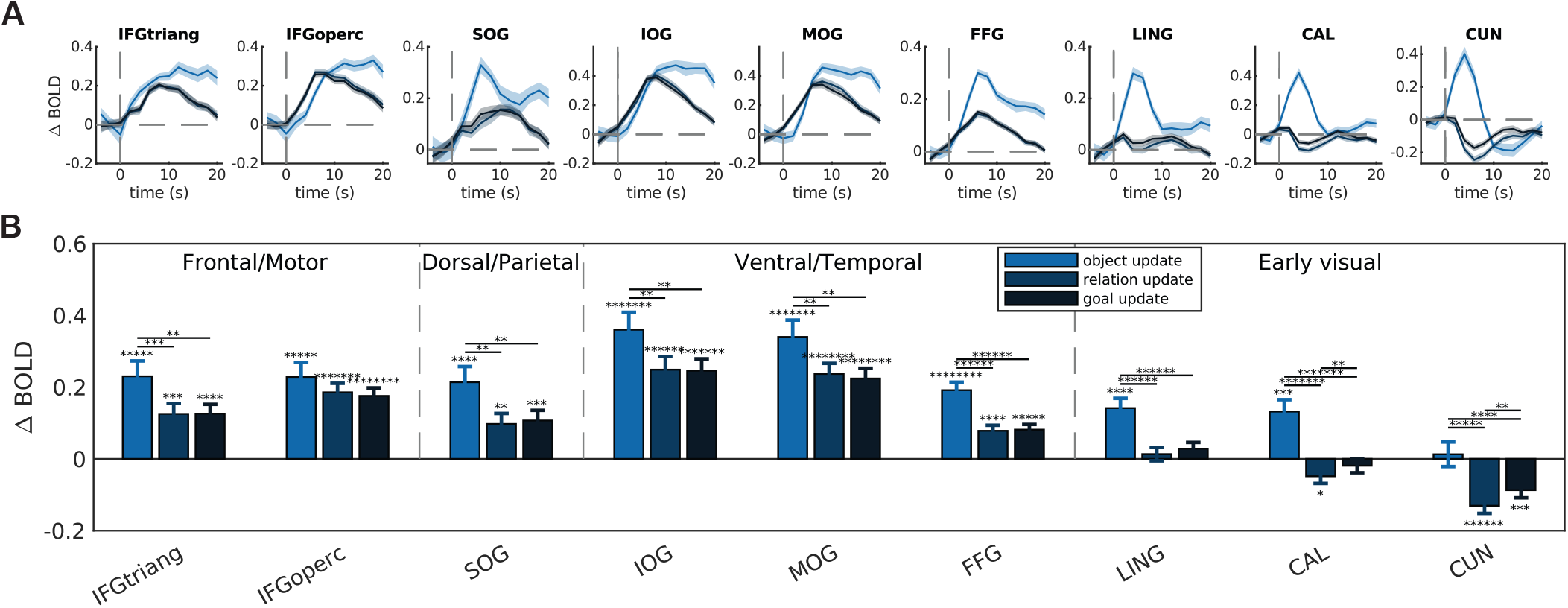
BOLD responses to individual theory component updates. Related to Figure 4C,D. A. Peri-event time histograms showing the average change in BOLD signal following individual theory component updates. Notation as in Figure 4C. B. Change in BOLD signal from A averaged over 20 s following corresponding event. Notation as in Figure 4D.

**Figure S6:**
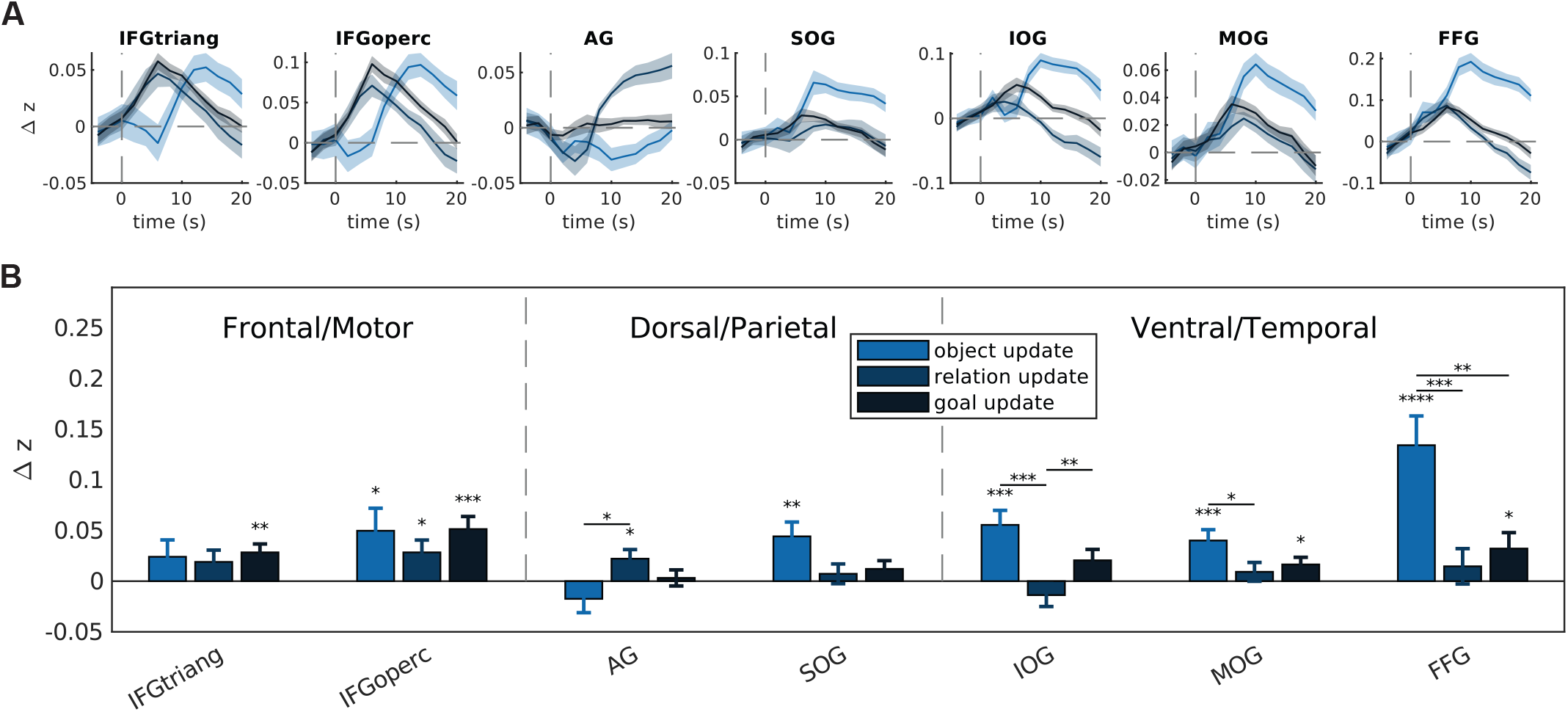
Thery component activation following corresponding component updates. Related to Figure 6B,C. A. PETHs showing the average change in predictivity score following individual theory component updates. Notation as in Figure 6B. B. Change in predictivity score from A averaged over 20 s following corresponding event. Notation as in Figure 6C.

**Table S1:** Theory representation clusters from encoding model t-map. Related to Figure 3. BA, Brodmann area. T-stat, t-statistic in peak voxel in cluster. MNI, Montreal Neurological Institute coordinates of peak voxel in cluster.

| Sign | AAL3 region | BA | Extent | T-stat | MNI |
| --- | --- | --- | --- | --- | --- |
| Positive | Fusiform gyrus (L) | 37 | 6655 | 9.035 | -36 -58 -14 |
|  | Middle occipital gyrus (L) | 19 | 6655 | 7.869 | -32 -86 16 |
|  | Superior occipital gyrus (L) | 17 | 6655 | 7.374 | -8 -100 12 |
|  | Cuneus (R) | 17 | 3088 | 7.004 | 18 -98 8 |
|  | Fusiform gyrus (R) | 37 | 3088 | 6.109 | 38 -52 -18 |
|  | Middle occipital gyrus (R) | 19 | 3088 | 5.231 | 38 -88 2 |
|  | IFG pars triangularis (L) | 48 | 1534 | 6.998 | -36 24 22 |
|  | IFG pars triangularis (L) | 45 | 1534 | 6.497 | -52 34 12 |
|  | IFG pars triangularis (L) |  | 1534 | 4.910 | -26 48 34 |
|  | Middle temporal gyrus (L) | 22 | 422 | 5.417 | -56 -32 8 |
|  | Middle temporal gyrus (L) | 22 | 422 | 4.141 | -32 -38 8 |
|  | Rolandic operculum (R) | 48 | 452 | 5.238 | 40 -26 22 |
|  | Supramarginal gyrus (R) | 2 | 452 | 5.072 | 58 -30 30 |
|  | Supramarginal gyrus (R) | 3 | 452 | 4.615 | 36 -22 42 |
|  | IFG pars opercularis (R) | 44 | 384 | 5.207 | 42 8 30 |
|  | IFG pars triangularis (R) | 48 | 384 | 4.892 | 42 26 20 |
|  | Precentral gyrus (R) |  | 273 | 5.160 | 54 -12 54 |
|  | Precentral gyrus (R) | 6 | 273 | 4.559 | 34 -10 48 |
|  | Cerebellum (R) |  | 224 | 5.041 | 14 -82 -28 |
|  | Cerebellum (R) |  | 224 | 3.955 | 38 -74 -40 |
|  | Supplementary motor area (L) | 6 | 260 | 4.984 | -8 18 64 |
|  | Middle frontal gyrus (L) | 9 | 260 | 4.682 | -32 14 54 |
|  | Precuneus (R) | 5 | 288 | 4.840 | 8 -48 56 |
|  | Postcentral gyrus (R) | 3 | 288 | 4.756 | 28 -40 52 |
|  | Postcentral gyrus (R) | 1 | 288 | 3.613 | 28 -42 72 |
|  | Middle occipital gyrus (R) | 19 | 284 | 4.799 | 34 -66 36 |

**Table S2:**
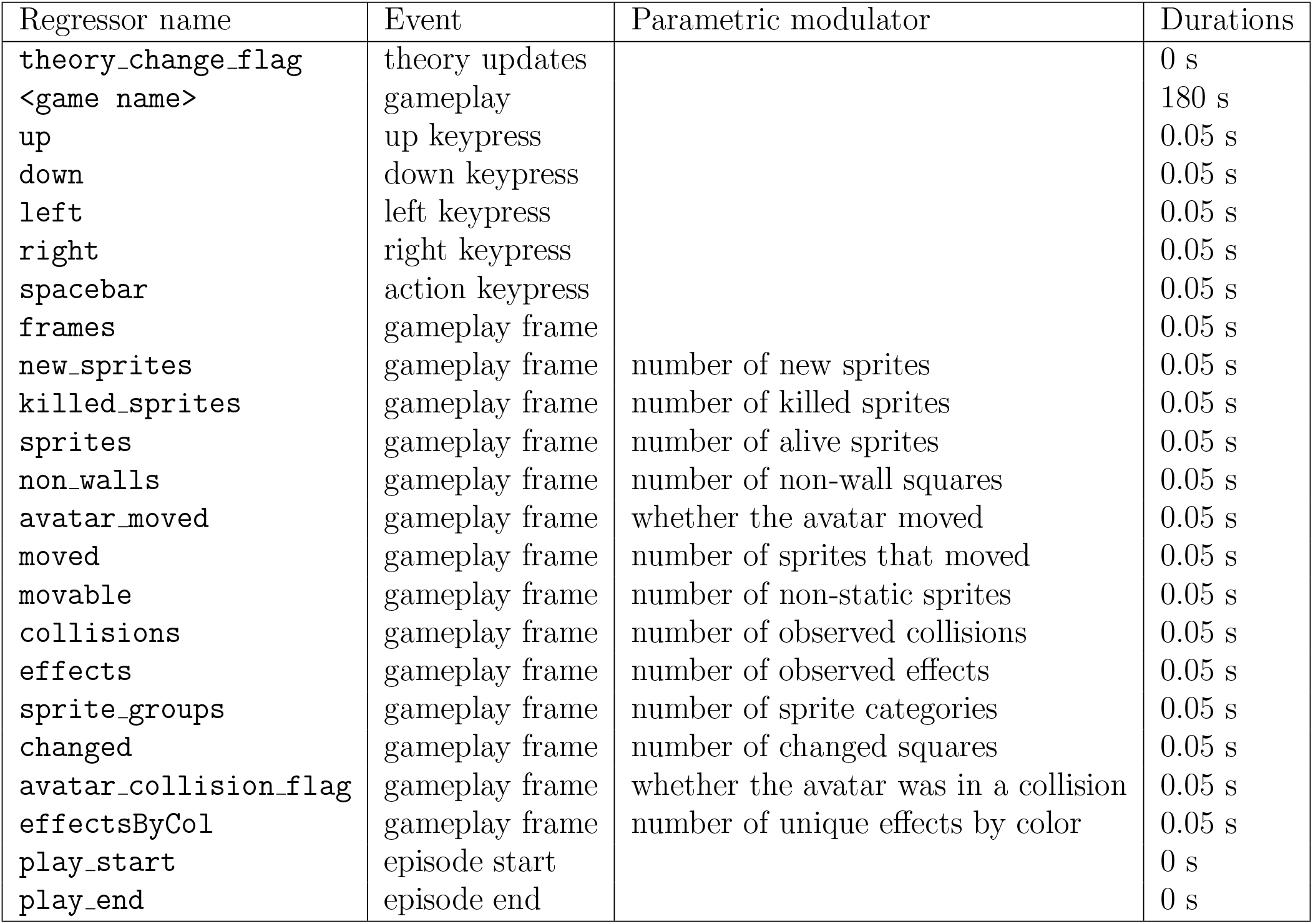
Theory update GLM regressors. Related to Figure 4.

**Table S3:**
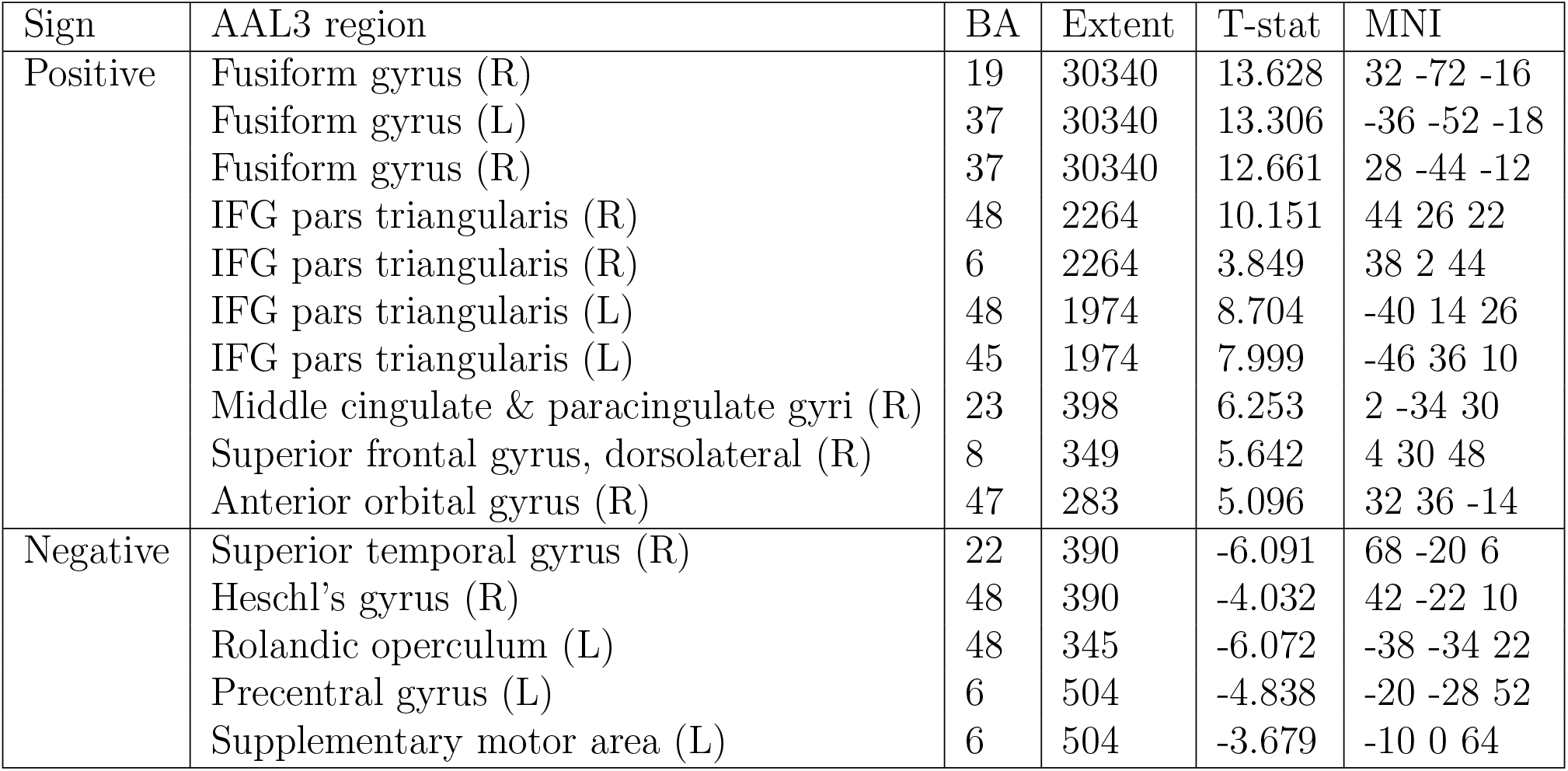
Theory updating clusters from GLM t-map. Related to Figure 4. Notation as in Table S1.

| Sign | AAL3 region | BA | Extent | T-stat | MNI |
| --- | --- | --- | --- | --- | --- |
| Positive | Fusiform gyrus (R) | 19 | 30340 | 13.628 | 32 -72 -16 |
|  | Fusiform gyrus (L) | 37 | 30340 | 13.306 | -36 -52 -18 |
|  | Fusiform gyrus (R) | 37 | 30340 | 12.661 | 28 -44 -12 |
|  | IFG pars triangularis (R) | 48 | 2264 | 10.151 | 44 26 22 |
|  | IFG pars triangularis (R) | 6 | 2264 | 3.849 | 38 2 44 |
|  | IFG pars triangularis (L) | 48 | 1974 | 8.704 | -40 14 26 |
|  | IFG pars triangularis (L) | 45 | 1974 | 7.999 | -46 36 10 |
|  | Middle cingulate & paracingulate gyri (R) | 23 | 398 | 6.253 | 2 -34 30 |
|  | Superior frontal gyrus, dorsolateral (R) | 8 | 349 | 5.642 | 4 30 48 |
|  | Anterior orbital gyrus (R) | 47 | 283 | 5.096 | 32 36 -14 |
| Negative | Superior temporal gyrus (R) | 22 | 390 | -6.091 | 68 -20 6 |
|  | Heschl's gyrus (R) | 48 | 390 | -4.032 | 42 -22 10 |
|  | Rolandic operculum (L) | 48 | 345 | -6.072 | -38 -34 22 |
|  | Precentral gyrus (L) | 6 | 504 | -4.838 | -20 -28 52 |
|  | Supplementary motor area (L) | 6 | 504 | -3.679 | -10 0 64 |

## Notes

### Competing Interest Statement

The authors have declared no competing interest.

